# Elucidating the Role of Cerebellar Nuclei Parvalbumin Activity on Adolescent Reversal Learning

**DOI:** 10.64898/2026.08.20.746009

**Authors:** Tristan T. Lyle, Aaron Berkley, Jessica L. Verpeut

**Affiliations:** Arizona State University, Department of Psychology, Tempe, AZ, 85287 USA

**Keywords:** cerebellum, cerebellar nuclei, parvalbumin, fiber photometry, mouse, perineuronal nets, chemogenetics, adolescent development

## Abstract

The cerebellar nuclei (CN) has demonstrated its influence on cognitive behavior via the cerebello-cortico circuit, yet the role of CN critical period mechanisms and how they may influence cognitive behavior, such as parvalbumin (PV) expressing interneurons enwrapped by perineuronal nets (PNNs), is still unclear. Therefore, we investigated the role of the lateral CN (LCN) PV cell calcium activity while animals performed a visual discrimination touchscreen cognitive task. All animals received the PV cell calcium indicator GCaMP6f at postnatal day 21 (P21). We targeted the LCN critical period by manipulating neural activity in male mice using the inhibitory Designer Receptors Exclusively Activated by Designer Drugs (DREADDs) from postnatal day 21 to 35 or by injecting an Hapln1-AAV vector to selectively target LCN PNN development. After animals completed the visual discrimination task, cerebellar tissue was collected for viral recovery and antibody staining for PNN components, Hapln1 and aggrecan. Results revealed DREADD animals showed improved reversal learning, an increase in calcium response to learning-related activity and altered PNN expression (Hapln1 and aggrecan). Hapln1 treated animals displayed a decrease in final day acquisition performance, lower reversal performance compared to DREADD groups, a decrease in reversal calcium learning-related activity, and an increase in PNN expression (Hapln1). Together, these data provide further evidence of LCN mechanisms associated with learning as well as the importance of understanding region-specific critical periods of plasticity.

**Highlights:**

● Inhibitory DREADDs improved reversal learning by increasing calcium responses in learning
● Hapln1 overexpression impaired learning and reversal performance
● Perineuronal nets in the LCN modulates neural activity and learning
● Parvalbumin positive neurons change activity patterns as a result of Hapln1 overexpression

## Introduction

The cerebellum has been regarded principally as a coordinator of movement, gait, and posture,^1^ yet emerging evidence suggests the cerebellum plays a pivotal role beyond the motor domain, contributing to non-motor functions (e.g., cognitive flexibility).^2–9^ Anatomical tracing, neuroimaging, and lesion data demonstrate closed-loop circuits between the cerebellar nuclei (CN) afferents to widespread prefrontal, parietal, and temporal regions, thus providing a structural basis for non-motor cerebellar functions. For example, afferents from dorsolateral prefrontal cortex map into lateral posterior cerebellar lobules (Crus I/II) via the pontine nuclei and lateral CN (LCN) efferents project back to the cortex mediated through the ventrolateral thalamus.^8,10–14^ These circuits underscore cognitive domains such as working memory, language, attention, and even social processes.^15^ The effect from damaging or interrupting these circuits reveal deficits in executive functions, affective regulation, and visuospatial reasoning.^16,17^ As research continues to elucidate cerebellar involvement in non-motor functions, the evidence can aid in cerebellar dysfunction observed in cerebellar physical trauma,^17^ neurodevelopmental disorders,^6^ and even addiction.^18,19^

The LCN is functionally connected to the prefrontal cortex region during cognitive and social tasks, and in autism spectrum disorder (ASD), there are changes in functionality between these regions.^20,21^ Moreover, an imbalance between excitatory and inhibitory activity ratios has been found in patients with ASD, which has been hypothesized to be a result of precocious closure of developmental critical periods.^22^ In other words, a possible mechanism contributing to decreased functional connectivity between the cerebellum and prefrontal cortex may be driven by an early onset of inhibitory interneurons. For example, a specific subset of GABAergic interneurons express the calcium binding protein parvalbumin (PV) and have been shown to develop during postnatal life in anticipation of critical periods.^23,24^ PV-cells have unique properties that regulate pyramidal neuron output, such as high firing rates, narrow action-potentials, and reliable GABA release.^25–27^ PV-cells target perisomatic and axonal regions of principal cells enabling inhibitory control of principal cell output and are found to regulate excitatory and inhibitory balances within the hippocampus, auditory cortex, and prefrontal cortex.^28–30^ However, research seeking to understand LCN PV interneurons during postnatal critical periods and how they relate to cognitive behavior is lacking.

Understanding LCN mechanisms that emerge during critical periods is needed to better understand how cerebellar output shapes the cerebello-cortico circuit. Perineuronal nets (PNNs) are a specialized extracellular matrix (ECM) that emerge during the closure of critical period plasticity. PNNs principally envelop the soma and perisynaptic space of PV interneurons and have been found to play a critical modulatory role in synaptic plasticity, calcium dynamics, and flexible learning.^2,31,32^ PNNs are composed of a hyaluronan backbone, chondroitin sulfate proteoglycans (CSPGs), hyaluronan link proteins (Haplns) and Tenascin-R (TnR), that form a lattice-like structure that influence ion diffusion, stability of synaptic contacts and limit structural plasticity. Reopening critical periods within the CN through enzymatic degradation of PNNs (e.g., ChABC) not only changes the intrinsic excitability, membrane capacitance, and temporal firing of PV cells, but also facilitates learning commensurate with juvenile plasticity.^33^ Moreover, genetic knockout or disruption of Haplns, TnR, or CSPGs (e.g., aggrecan) can severely disrupt PNN composition, function, stability, and behavior.^32^ For example, among the three Hapln proteins abundant in the brain, Hapln1/Crt1 knockout mice long-term object recognition memory is prolonged similarly to PNN degraded animals, providing evidence that PNNs may regulate recognition memory.^34^ Moreover, deleting the aggrecan gene, the most abundant CSPG within PNN populations,^35^ prolonged recognition memory in mice 24-h after object exposure compared to controls.^36^ However, it is still unclear how Hapln1 and aggrecan are related to the LCN mediated cognitive behavior during critical periods.

In the current study, we hypothesized that typical maturation of the LCN neuronal population and PNN components are influential on cognitive-related behaviors. To investigate associative functions of critical period LCN neuronal development and cognitive behavior, we used chemogenetics to reduce LCN activity, a viral vector approach to selectively increase PNN component Hapln1, and a GCaMP6f viral vector to selectively record calcium in PV expressing interneurons in male mice. In adolescence, a visual discrimination task was used to evaluate changes in acquisition and reversal learning while LCN PV calcium activity was recorded. Postmortem LCN PNN components were examined to understand the relationships between LCN plasticity and cognitive behavior. Our findings displayed an improvement in reversal learning and increased calcium during error-related responses during reversal in LCN DREADD inhibited mice. Overexpressing Hapln1 using a Hapln1 adenoassociated virus, had specific impacts on late learning and reduced error-related calcium activity during reversal. These studies provide a foundation for exploring LCN critical period plasticity and relationships to cognitive behavior.

## Materials and Methods

### Mice

Male (n = 19, Table 1) C57BL/6J mice (JAX stock #000664, https://www.jax.org/strain/000664) used in this study were bred in-house in monogamous breeding pairs. We used male mice specifically as there is a higher prevalence in males within ASD populations ^37^ and there is a lack of evidence regarding cerebellar hormone specific changes in learning throughout the ovulation cycle.^38,39^ All mice were housed in ventilated top cages (Allentown) and received environmental enrichment, including paper nesting squares. Mice were provided Teklad Global Rodent Diets (Teklad Diets) and water ad libitum. Cages were changed every 2 weeks, and animals were housed in groups of 4−5 mice in reverse light cycle (12 h light/dark cycle) rooms to maximize normal nocturnal activity, as behavior testing occurred during the day. All mice were handled and housed in accordance with the guidelines of the Institutional Animal Care and Use Committee at Arizona State University (protocol #24-2052R).

**Table 1.**
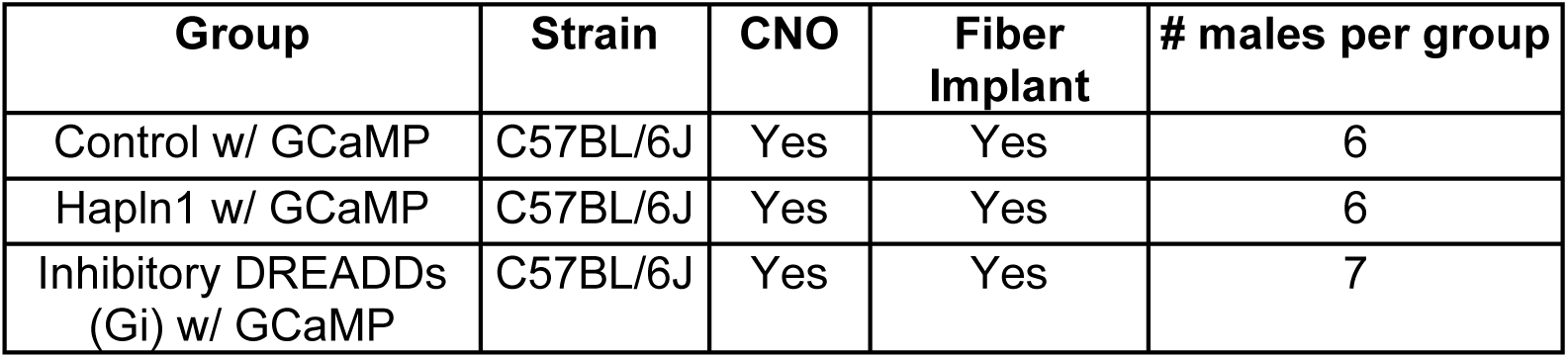
Experimental groups (Control, Hapln1, Gi) and designated groups receiving fiber implant and CNO in drinking water. All mice were male C57BL/6J.

| **Group** | **Strain** | **CNO** | **Fiber Implant** | **# males per group** |
| --- | --- | --- | --- | --- |
| Control w/ GCaMP | C57BL/6J | Yes | Yes | 6 |
| Hapln1 w/ GCaMP | C57BL/6J | Yes | Yes | 6 |
| Inhibitory DREADDs (Gi) w/ GCaMP | C57BL/6J | Yes | Yes | 7 |

### CN Manipulation Using Viral Vectors

Animals received surgery at P21 for transduction of GCaMP adeno-associated virus (AAV) and either inhibitory (Gi) Designer Receptors Exclusively Activated by Designer Drugs (DREADDs) or PNN component Hapln1-AAV in the LCN. Injection of Gi-coupled AAV carrying the sequence for the inhibitory DREADD under a human synapsin-1 promoter with a mCherry reporter (AAV2-hSyn-hM4D(Gi)-mCherry was a gift from Bryan Roth, Addgene viral prep # 50475-AAV2; http://n2t.net/addgene:50475; RRID:Addgene_50475; 1E+13 vg/ml) to drive expression of Gi in the LCN. Injection of Hapln1-AAV carrying the sequence for Hapln-1 under a human synapsin-1 promoter with a cfGFP2 reporter (AAVDJ-hSyn-Hapln1-cfGFP2 plasmid was a gift from Alexander Dityatev, Addgene plasmid # 200777; http://n2t.net/addgene:200777; RRID:Addgene_200777; recombinant AAV (DJ) with Hapln1-cfGFP2, 1E+13 vg was purchased from Virovek, Hayward, CA) to increase expression of Hapln1 in the LCN. All animals were also injected with GCaMP AAV carrying the sequence for GCaMP6f calcium sensor under a S5E2 promoter to target PV interneurons with an GFP reporter (pAAV-S5E2-GCaMP6f was a gift from Jordane Dimidschstein, Addgene viral prep # 135632-AAV9 ; http://n2t.net/addgene:135632; RRID:Addgene_135632; 1E+13 vg/ml), to drive GCaMP6f expression in LCN PV interneurons. All mice (n=19) had access to the DREADD activator, clozapine-*N*-oxide (CNO; 10 mg/kg, NIMH Chemical Synthesis and Drug Supply Program) in drinking water, similar to our previous study,^2^ from P21 to P35 to selectively activate the inhibitory hM4D (Gi) muscarinic receptor. Control animals only received CNO in their drinking water and the GCaMP AAV (n=6).

### Stereotaxic Surgery

Procedures were performed similar to Lyle and Verpeut 2025.^2^ Briefly, mice were first anesthetized with 5% isoflurane and subsequently maintained with 2–3% isoflurane. Body temperature and respiration was monitored every 5 minutes. Mice were secured in a stereotaxic setup (David Kopf Instruments, Tujunga, CA) via ear bars and kept warm on a heating pad (RightTemp Jr, Kent Scientific). Animals received the osmotic diuretic drug, mannitol (15% D-mannitol in DPBS, 0.2ml, intraperitoneal, Sigma-Aldrich, Cat #M4125). Hair was removed from the scalp and cleaned. An incision was made along the lambdoid suture and cut posteriorly until it reached the cerebellum. Bilateral craniotomies were performed over both dentate nuclei with a 0.5 mm micro-drill burr (Micromotor Drill, Stoelting, cat #51449). The DREADD (250 nL) or Hapln1 (250 nL), and GCaMP virus (250 nL) was administered via hamilton syringe (700 series, Fisher Scientific, cat #87930) using the coordinates from the Paxinos and Franklin mouse brain atlas targeting the dentate nuclei in the cerebellum: (AP) = -5.80 mm, (ML) = 2.3 mm, (DV) = -1.5. The craniotomy was sealed with Kwik-Sil before closing the incision. Wounds were sutured (5-0, Ethicon) closed and tissue glue (Vetbond) applied on top of the sutures. Mice were then injected subcutaneously with Rimadyl (0.2 mL) and intraperitoneally with 15% D-mannitol (0.5 mL) and were monitored for full ambulatory recovery prior to being returned to their home cages. For fiber photometry experiments, mice were unilaterally implanted with an 1 mm optical fiber (0.66 NA, 225/245 μm diameter) after animals completed the shaping phase of the VD task (∼10 d) in the LCN. The fiber implant was fixed in place by gluing it to the skull with C&B-Metabond (Parkell Inc., Edgewood, NY, USA) according to the manufacturer’s instructions. All recordings were made using the Multi-Wavelength Photometry System from Plexon. PlexBright optogenetic stimulation system custom length fiber stub implants (ferrule magnetic-tipped patch cable; 225/245um diameter fiber; flat cleaved; magnetic LC ferrule; 1mm length) were used for neural recordings at 465 nm for GCaMP6f. Post-sacrifice all fiber implants were verified for proper location through histology to locate the cannulation lesion.

### Visual Discrimination and Touchscreen Task

Freely moving animals were conditioned to visually discriminate between two stimuli by using a modular touch screen operant chamber (Med Associates), which was operated using commercial software (K-Limbic Touch Screen Software) and placed in a noise-attenuating chamber, similar to Lyle and Verpeut 2025.^2^ The following preshaping stages were implemented: 1) Animals habituated to the experimental environment for single 30 min sessions. 2) Mice were trained to initiate a trial by moving their head into the reward area, crossing the infrared beam, and then touching the screen regardless of the side. The criteria to move on from Stage 2 was a 60% accurate response. 3) Mice were trained to touch only the target stimulus within a 30 s onset, the trials time out if no response occurred or an incorrect response occurred, and the next intertrial interval starts. Correct responses were reinforced with 15% sweetened condensed milk. Mice were required to reach performance criteria of 80% or higher accuracy to move onto visual discrimination with sessions lasting 30 min per day. During visual discrimination testing, the mice were presented with two visual stimuli and were only reinforced for touching the target stimulus. An error in selecting the wrong stimulus or a failure to respond within 30 s ended the trial without a reward. The mice performed visual discrimination for 5 d and then moved onto the reversal stage. During reversal, the contingency switched where reinforcement was contingent upon touching the previous distractor stimulus. Animals were tested for an additional 5 d on reversal.

Trial initiation, response, and reward collection latencies were calculated by taking the average median using the median function in Microsoft Excel for each mouse and calculating the cumulative sum, respectively. Reward collection latency medians were only collected for trials in which an animal responded correctly, and the reward was available.

### Perfusion

Mice were anesthetized with 5% isoflurane and then subcutaneously injected with 0.2 ml of Euthasol (390 mg pentobarbital sodium, 50 mg phenytoin sodium, Virbac). Subsequently, blood was replaced with ∼30 ml phosphate-buffered saline (PBS) and then ∼50 ml 4% paraformaldehyde (PFA) over ∼10 min. Next, the brains were removed and postfixed in 4% PFA overnight. After 24 h, the tissue was transferred into ∼20% sucrose and kept at 4°C until sectioning.

### Tissue Collection and Immunostaining

Brain tissue was cut into coronal sections (40 μm) using a cryostat (Leica CM1860), and free-floating sections were placed into well plates containing PBS. Briefly, tissue was washed (three times for 10 min) in PBS and then blocked with 10% donkey serum, 0.5% Triton X-100, and PBS for 1 h at room temperature. Next, tissue was stained with mouse anti-aggrecan (1:1000, Millipore Sigma, catalog #MABT84) or rabbit anti-Hapln1 (1:250, Thermo Fisher Scientific, catalog #MA5-32100) in 2% donkey serum, 0.4% Triton X-100, and PBS for 48–72 h gently rocking at 4°C. Tissue was then washed (three times for 10 min) with PBS and stained with donkey anti-mouse Alexa Fluor 647 (1:500, Thermo Fisher Scientific, catalog #A31571) or donkey anti-rabbit Alexa Fluor 647 (1:1,000, Thermo Fisher Scientific, catalog #A31573) and Dapi (1:1000, Thermo Fisher Scientific, catalog #62248) in 2% donkey serum, 0.4% Triton X-100, and PBS. Tissue was then placed on glass slides (25 × 75 cm) and mounted using ProLong Diamond (Thermo Fisher Scientific, catalog #P36961).

### Microscopy and Image Analysis

All analysis was performed using Zen Blue, Image J, or QPath and analyzed using custom Python and R scripts. One Hapln1 mouse was removed from analysis due to poor tissue quality post-extraction and groups were balanced using only best tissue quality. Mouse brain tissue was imaged on an epifluorescent Zeiss Imager M2 using multiple fluorescent filter sets and a mechanical stage for *z*-stacks of DAPI, Hapln1, aggrecan, GFP, and mCherry with a set of objectives (20x), multiple fluorescent filter sets, and a mechanical stage for *z*-stack imaging. Epifluorescent images of at least three anatomically matched sections that include the LCN were quantified for viral expression (mCherry; GFP) in the targeted region, location, and PNN components Hapln1 and aggrecan pixel intensities.

LCN Hapln1 and aggrecan images taken on the epifluorescent Zeiss Imager M2 had a constant set of imaging parameters: Hapln1 secondary antibody LED channel power (50%) and exposure time (14 ms), aggrecan secondary antibody LED channel power (50%) and exposure time (14 ms), mCherry LED channel power (50%) and exposure time (14 ms), GFP LED channel power (50%) and exposure time (14 ms) and DAPI LED power (50%) and exposure time (6 ms). At 20× objective, a 10 μm *z*-stack was performed for each image and then max-projected using the Zeiss Imager M2 software similar to Lyle and Verpeut 2025.^2^ LCN Hapln1 and aggrecan mean pixel intensities were gathered by tracing each cell using the Qupath ^40^. We then calculated the average mean pixel intensity of a section by dividing the traced cell pixel intensity values by the total number of cells traced within the respective section. Images acquired on the Zeiss LSM 710 confocal at 40x used a Plan-Apochromat 40×/0.95 Korr objective. DAPI, GFP/GCaMP, and Alexa Fluor 647 were excited using 405, 488, and 633 nm laser lines, respectively. Images were acquired at 2196 × 2196 pixel resolution with a pixel size of 0.0968 × 0.0968 µm and z-step intervals of 0.992 µm. To calculate the percentage of viral DREADDs mCherry (Gi; n=4) and GCaMP-GFP (Gi, n=4; Hapln1, n=4; Control, n=4) reporters within each CN, we analyzed three coronal sections containing the CN from each mouse, respectively. We first measured the amount of viral reporter (mCherry; GFP) spread within each CN by circling the viral reporter (mCherry; GFP) using the measurement tool in Zeiss Zen software. Then, each CN was circled to calculate the total area per CN. Each viral reporter area measurement (microns squared) was divided by the total area of each CN. Additionally, AAV spread into the vestibular nuclei was performed the same as the mCherry and GFP spread within each CN in Gi (n=3) and Hapln1 (n=3) animals by analyzing three sections per mouse and bilateral vestibular nuclei regions.

## Data Analysis

Five parameters were collected on a session-by-session basis: completed trials, correct and incorrect performance, trial initiation latency, response latency, collection latency, and calcium recordings. All statistical analyses were conducted using RStudio (version 4.09.1). One-way ANOVAs were conducted to assess significant group differences in performance, completed trials, and latencies and post hoc Tukey’s HSD were conducted to examine significant pairwise comparisons within these parameters for both acquisition and reversal. At least 5 mice per group and 3-5 sections per mouse were collected for post-reversal LCN Hapln1 and aggrecan pixel intensities. Pixel intensities were averaged across all images collected from individual mice respectively. The average mean pixel intensity per section was calculated through the Microsoft Excel formula. All LCN Hapln1 and aggrecan averages were then analyzed using a linear regression model and subsequent One-way ANOVAs to further assess LCN Hapln1 and aggrecan variance between groups and post hoc Tukey’s HSD pairwise-comparisons tests unless indicated otherwise. Plexon fiber photometry csv files were exported and GCaMP and isosbestic signals were transformed to detrended/corrected signals to account for photobleaching/photoswitching ^3,41^ using custom R and Matlab scripts. For both acquisition and reversal each session GCaMP and isosbestic signal was transformed by applying an exponential model as a decreasing function per animal and session. Z-score and Area-under-the-curve (AUC) values were calculated using pMAT, an open source software for fiber photometry analysis,^42^ from 1 second prior to an animal engaging in a response (pre-event choice) to 1 second after the response (post-event choice). Area-under-the-curve (AUC) values were analyzed using linear mixed-effects models (LMMs; lme4 package, version 1.1-29) to account for the repeated-measures structure of the data, with mouse included as a random intercept. Fixed effects included Epoch (pre-vs post-event), Group, Day, and all interaction terms. To assess the statistical significance of main effects and interactions among fixed factors, a Type III analysis of variance (ANOVA) with Satterthwaite-approximated degrees of freedom (lmerTest package) was conducted on the fitted LMM. This approach allows for testing of fixed effects while retaining the mixed-effects structure of the model. Post hoc analyses and simple slopes were calculated when appropriate using estimated marginal means (emmeans package, version 1.7.4-1), with p-values adjusted for multiple comparisons.

## Results

### Experimental design

To elucidate the impact of the CN on cognitive behavior during a period of neural circuit development, we used chemogenetics to selectively inhibit cells in the CN from P21 to P35 or manipulated ECM structure by overexpressing Hapln1. All mice also received GCaMP6f for calcium measurements prior to cannulation surgery and behavior testing with neural recordings (**Figure 1a**). The inhibitory DREADDs or Hapln1 vector, was expressed in bilateral LCN cells using an adeno-associated virus (AAV) with a human synapsin 1 gene (hSyn) promoter ^43^ at P21 along with GCaMP6f-S5E2-AAV to express GCaMP6f in parvalbumin neurons (**Figure 1b**). All mice received the DREADD activator, clozapine-*N-*oxide (CNO; 10 mg/kg) in the drinking water from P21 to P35. At P36, CNO water was removed and replaced with typical drinking water for the rest of the experiment. After shaping, animals received surgery for the fiber photometry recording cannula (Figure 1c) and recovered for 1 week. After recovery, animals performed acquisition and reversal while GCaMP signal was recorded across all sessions for each mouse respectively (Figure d). After completing reversal, animals were perfused and cerebellar tissue was collected for viral recovery, cannula placement verification, and immunohistochemistry. It is critical to analyze the recovered injection site to control for any possible expression difference of the adenoassociated virus between individuals. To validate the expression of the DREADD and GCaMP viruses within our target and surrounding regions we analyzed DREADD expression analysis within the CN using the mCherry-positive voxel analysis found viral spread into ∼56% of the anterior LCN, 23% of the interposed, 16% in the fastigial, and 10% in the vestigial nuclei (**Figure 1e-f**). We performed the same voxel analysis for GCaMP expression and found spread within ∼55% of the LCN, 39% of the interposed, 24% of the fastigial, and 17% of the vestigial nuclei (**Figure 1f**).

**Figure 1.**
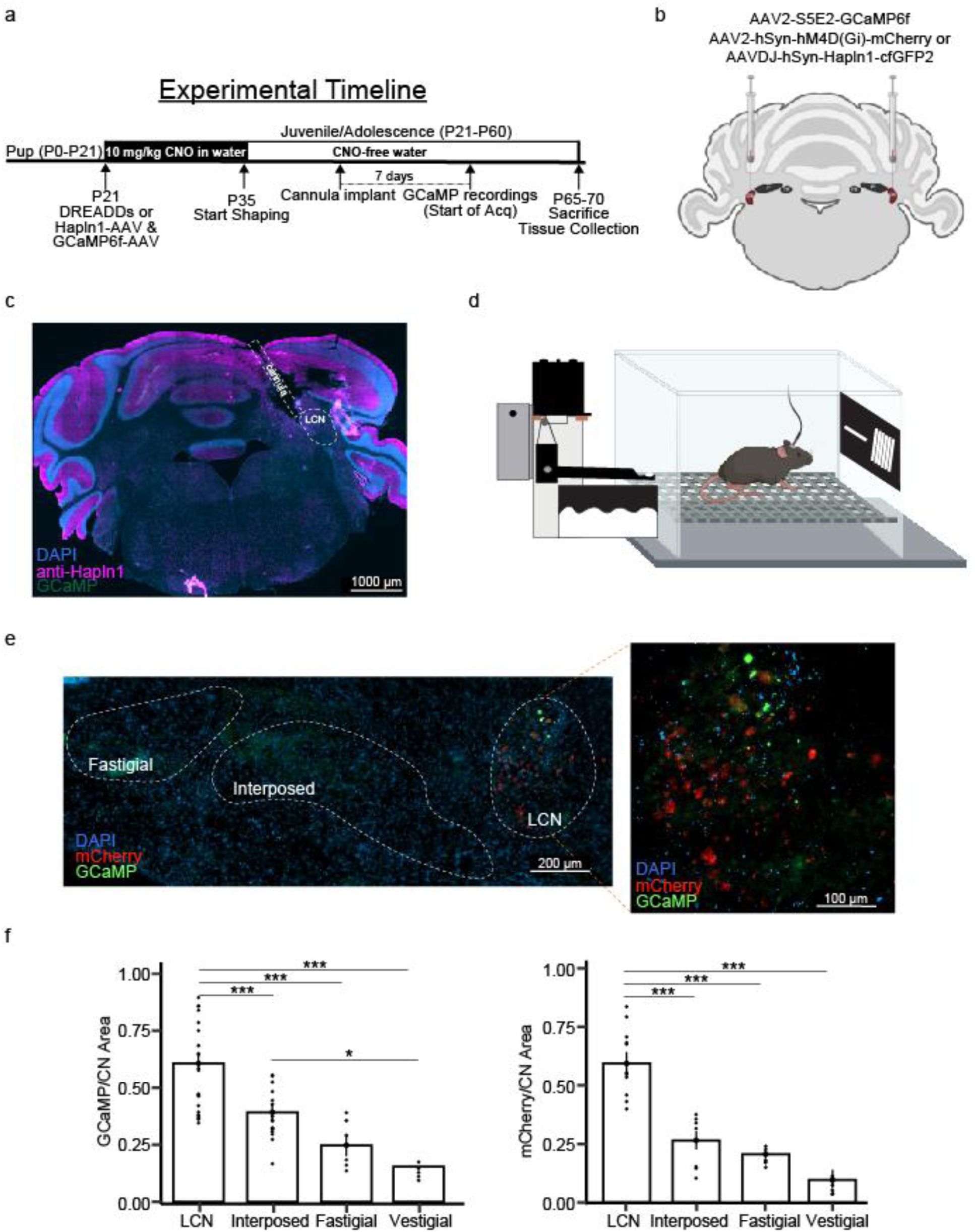
Experimental approach. (A.) Experimental timeline indicating the bilateral intracranial infusion of the inhibitory AAV-DREADD (Gi) or AAV-Hapln1 and AAV-S5E2-GCaMP6f into the lateral cerebellar nuclei (LCN) at P21. DREADD ligand clozopine-*N-oxide* (CNO; 10 mg/kg) was provided in drinking water for 2 weeks and replaced with normal water at P35 when animals started the visual discrimination task. After completing the shaping stage, all animals received the fiber photometry implant into the left LCN and recovered in their home cage for 1 week. After recovery, animals performed the acquisition phase for 5 days and then performed the reversal phase for 5 days. GCaMP signals were recorded across all sessions during acquisition and reversal for each animal respectively. After completing reversal, animals were perfused and cerebellar tissue was collected for viral recovery and immunohistochemistry of perineuronal nets components (B.) Cartoon illustration of a cerebellar coronal section displaying the CN and infusion of bilateral GCaMP and DREADDS or the Hapln1 viral vector. Created with BioRender.com, adapted from.^2^ (C.) Example image of a coronal section of the cerebellum indicating where the photometry fiber was inserted into the CN taken at 20x objective (DAPI: blue, anti-Hapln1: magenta, GCaMP: green). (D.) Cartoon example of a mouse performing the visual discrimination task while recording GCaMP activity. Created with *BioRender*.com. (E.) Example cerebellar nuclei image (DAPI: blue, anti-Hapln1: magenta, GCaMP: green) imaged at (left) 20x and (right) the LCN specifically at 40x. (F.) Fraction of the GCaMP reporter, GFP in animals receiving both GCaMP and DREADDs (mCherry). There was a significant difference between regions analyzed (F(3,60)=8.032, p=0.00014, ANOVA with the LCN having the highest level of GCaMP expression compared to the interposed (p=0.00049, Tukey), fastigial (p=0.00036, Tukey), and vestigial (p=0.00043, Tukey) nuclei. Additionally, the interposed had greater GCaMP expression compared to the vestigial nucleus (p=0.035, Tukey). The DREADD reporter, mCherry, was significantly different across regions analyzed (F(3,21)=13.41, p=0.00004, ANOVA) with the highest amount found in the LCN compared to the interposed (p=0.0002, Tukey), fastigial (p=0.0001, Tukey), and vestigial (p=0.00019, Tukey) nuclei. Data represents mean ± SEM, *p < 0.05, **p < 0.01, ***p < 0.001.

### Pairwise Discrimination Task to Study Cognition

Here, we utilized the touchscreen pairwise visual discrimination task, a translational tool from mice-primates-humans that allows for examination of learning and flexible cognition ^2,44^. We trained animals in the visual discrimination task using an automated touchscreen-based pairwise discrimination protocol, similar to Lyle and Verpeut 2025.^2^ Briefly, mice were shaped to self-initiate a trial by breaking an infrared beam at the reward port, then touch the touchscreen using either their nose or paw, and finally touch the screen when an image was presented. After shaping, animals underwent the acquisition phase where mice were rewarded for touching the target stimulus (e.g., vertical white bars) with 15% sweetened and condensed milk. Animals were tested for 30 min per day for 5 d to acquire the visual discrimination contingency (acquisition). After 5 d, the contingency was reversed (i.e., nontarget stimulus was now reinforced) to examine cognitive flexibility (reversal) for an additional 30 min per day for 5 d. The figures were randomized in their presentation across both screens. All animals were sacked and perfused for analysis of biological markers to explore the relationship with reversal performance.

First, we analyzed the shaping phase of our paradigm (Stages 1–4) by calculating each the shaping index. Shaping index was calculated by dividing the cumulative trials completed by the total number of days per stage. We then performed a two-way ANOVA with group and stage as factors to explore potential differences in the amount of trials completed in each stage (**Figure 2**). Our results showed a main effect of group (F(2,70)=4.58, p=0.01, ANOVA), and Hapln1 animals, regardless of stage, required more trials to complete shaping compared to Gi animals (p=0.01, Tukey). We found no main effect of stage (F(1,70)=0.04, p=0.84, ANOVA) or interaction between factors Group and Stage (F(2,70)=0.09, p=0.91, ANOVA) predicting the proportion of completed trials. Overall, these results show that Hapln1 animals required more trials overall to complete the shaping phase of the learning task.

**Figure 2.**
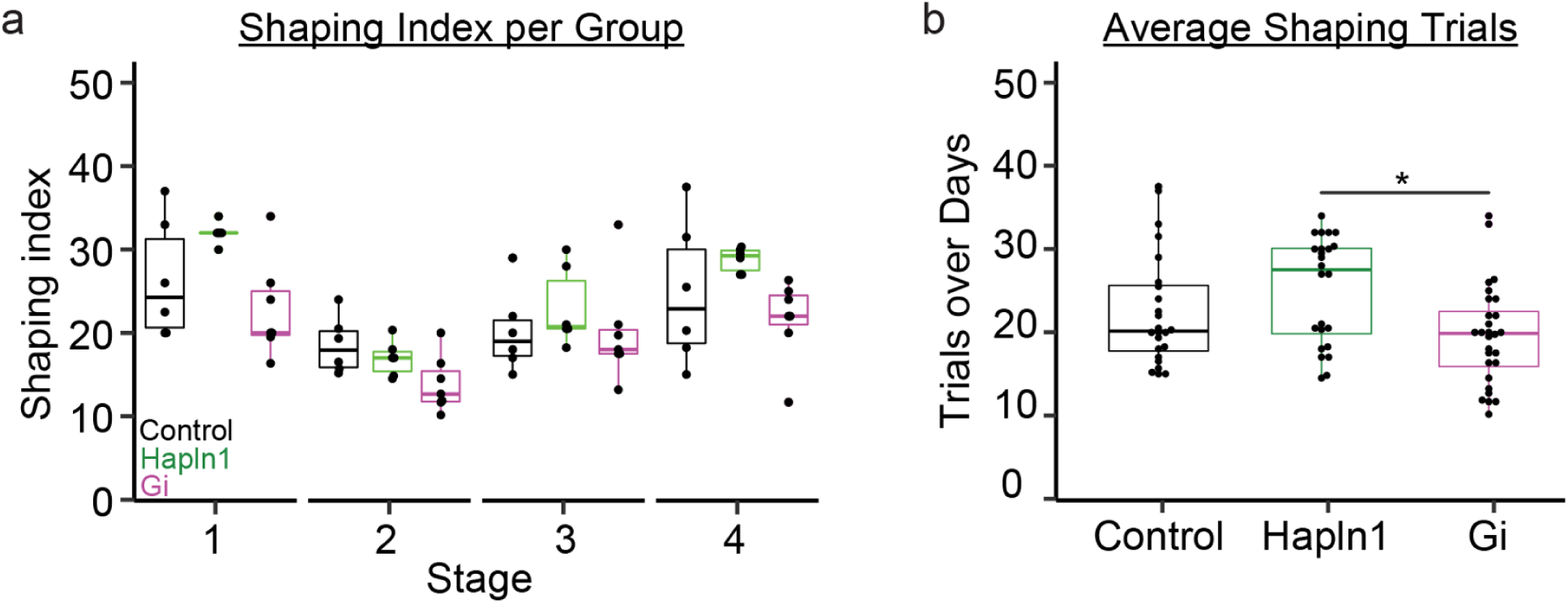
Learning stages of the visual discrimination task. (A.) Shaping index was calculated by dividing the cumulative trials completed by the total number of days per stage. There were no stage specific factors found to influence the number of trials completed overall (F(1,70)=0.043, p=0.84, ANOVA). (B.) Overall group differences were found (F(2,70) = 4.85, p = 0.01, ANOVA), driven specifically by Hapln1 animals (n = 6) requiring more trials to complete shaping compared with Gi (n = 7) groups (p = 0.01, Tukey). Data represents mean ± SEM, *p < 0.05

### Increasing Hapln1 Impacts Final Day of Acquisition

For acquisition performance analysis, we performed a logit transformation of correct choice percentage, similar to Lyle and Verpeut 2025.^2^ We first compared logits across acquisition to identify any potential differences in performance using a two-way ANOVA with Group and Day as factors. Our results revealed no Group differences in logits across all acquisition days (F(2,91)=0.95, p=0.39, ANOVA) or main effect for Day (F(1,91)=1.9, p=0.17). Additionally, logit slope analysis showed no differences between groups (F(2,16)=0.69, ANOVA). However, analysis of the last day of acquisition, revealed logit differences (F(2,16)=3.95, p=0.04, ANOVA), specifically between Hapln1 animals having lower performance compared to Gi animals (p=0.04, Tukey, **Figure 3a**). There were no differences in the final day of acquisition performance between Control and Gi animals (p=0.74, Tukey) or Hapln1 animals (p=0.16, Tukey). Our latency analysis revealed no group differences in trial initiation latency (F(2,16)=1.63, p=0.23, ANOVA), responding to the stimulus latency (F(2,16)=0.64, p=0.54, ANOVA), or collecting a reward latency (F(2,16)=0.73, p=0.49, ANOVA) (**Supplementary Figure 1**). Lastly, we explored whether the number of completed trials was related to acquisition performance and found that trials were not a significant predictor (p=0.64) of performance. Further exploration of trial completion revealed group differences in the cumulative sum (F(2,16)=4.983, p=0.02, ANOVA), specifically Hapln1 animals completing more trials than Controls (p=0.02, Tukey) but no different than Gi animals (p=0.11, Tukey) (**Figure 3b**). Together, these results suggest that DREADD manipulation did not alter acquisition. However, Hapln1-treated animals did complete more trials and displayed lower performance on the last day of acquisition.

**Figure 3.**
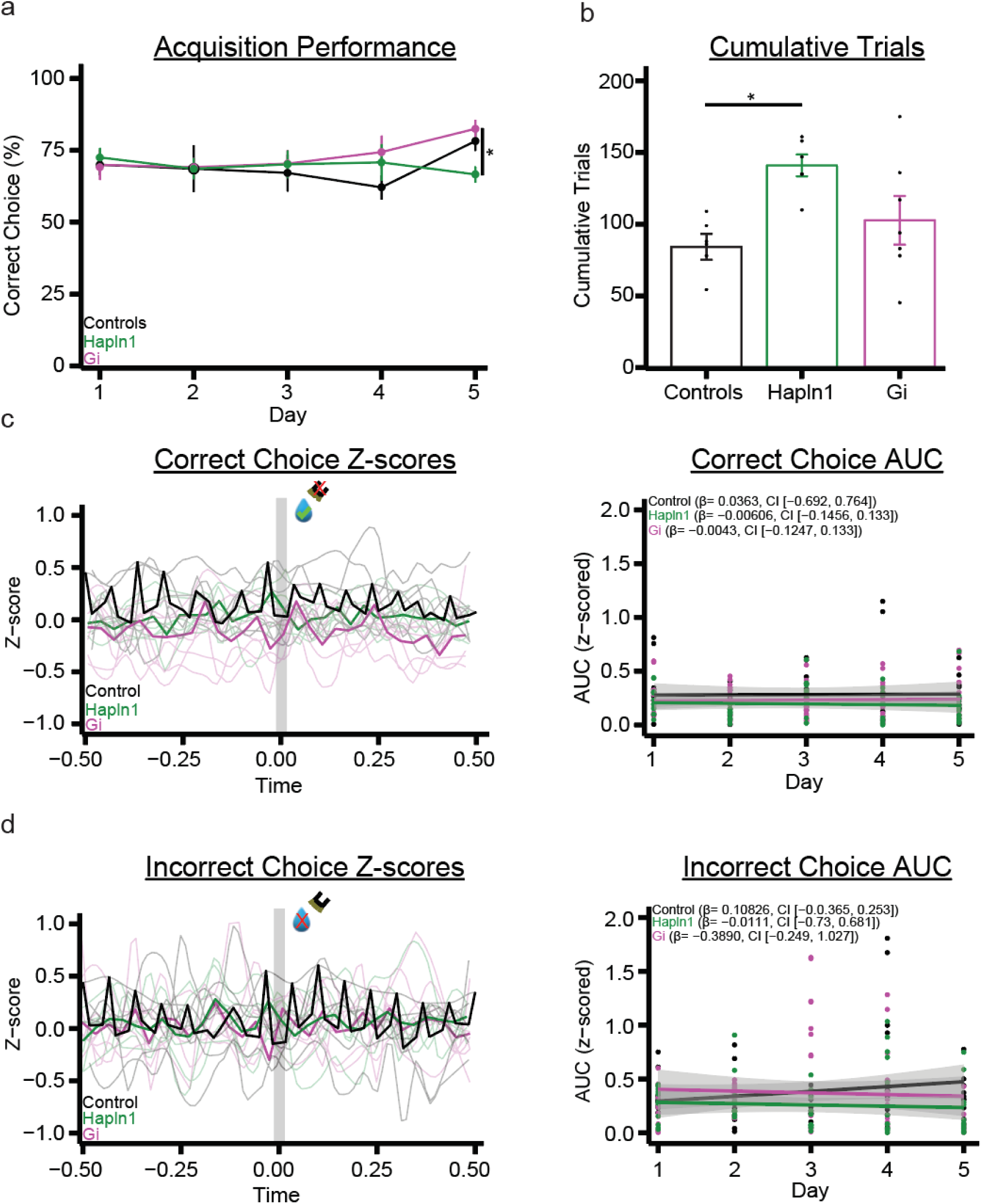
Increasing Hapln1 reduces final acquisition day without changing calcium activity. Control (black, n=6), Hapln1 (green, n=6), Gi (magenta, n=7). (A.) Correct choice (%) over 5 days of the task acquisition. A logit transformation of acquisition proportions of correct choice (%) was performed. A linear regression using the logit scores was then performed to calculate the slopes and analyzed using One-way ANOVAs with group as a factor. Mice showed no differences in logit slopes between groups during acquisition and reversal. Analysis of the final day of acquisition (day 5) logit scores revealed significant group differences (F(2, 16)=3.95, p=0.04), and Gi animals (n=7) displayed higher performance compared to Hapln1 (n=6) animals (p=0.036, Tukey). (B.) Comparison of group cumulative completed trials across acquisition (F(2,16)=4.983, p=0.02) revealed Hapln1 animals completed more trials overall compared to controls (p=0.02, Tukey). No differences were found between Hapln1 and Gi or control and Gi animals. (C.) (Left) Visualized GCaMP z-scores from 0.5 s before and 0.5 s after a correct choice occurred. Vertical grey bar indicates at timepoint zero when a response occurred and semitransparent lines are averaged correct choice z-scores per animal with bold lines indicating group averages. (Right) Area-under-the-curve (AUC) slope analysis for correct choice instances were calculated by first computing an average z-score for each correct choice occurrence for each animal and day respectively. AUC was calculated for each animal by using their average correct choice z-score for each day. Simple-slopes analysis of AUCs were then performed using AUCs marginal means trends. Simple Slope analysis revealed no significant changes in correct-choice calcium activity across acquisition for control (β= 0.0363, CI [−0.692, 0.764], Hapln1 (β= −0.00606, CI [−0.1456, 0.133]), or Gi (β= −0.0043, CI [−0.1247, 0.133]) animals. There were no differences in slopes between groups. (D.) (Left) Visualized GCaMP z-scores from 0.5 s before and 0.5 s after an incorrect choice was performed. Vertical grey bar indicates at timepoint zero when an incorrect choice occurred and semitransparent lines are averaged incorrect choice z-scores per animal with bold lines indicating group averages. AUC slope analysis for incorrect choice was the same as for correct choice, except calculated from incorrect choice occurrences. Analysis revealed no significant changes in incorrect-choice calcium activity across acquisition for controls (β= 0.10826, CI [−0.0.365, 0.253]), Hapln1 (β= −0.0111, CI [−0.73, 0.681]), or Gi (β= −0.3890, CI [−0.249, 1.027]) animals. Additionally, there were no differences in slopes between groups. Data represents mean ± SEM, *p < 0.05

### Acquisition Calcium Activity Remains Constant

To understand changes in local firing rate as a result of DREADD and Hapln1 manipulation and whether LCN PV cell population activity was related to learning, we analyzed changes in calcium using the genetically encoded calcium indicator (GCaMP6f) while animals performed the visual discrimination task. First, we calculated GCaMP z-scores from -1 s (pre-event) to 1 s after (post-event) a response occurred (correct or incorrect choice) for each animal per day respectively. Next, we calculated the area-under-the-curve (AUC) values for each animal per day based on correct and incorrect choice. We then performed linear mixed-effects models (LLMs) to predict AUC values with fixed effects Epoch (−1 to 0 s, pre-event; 0 to 1 s, post-event), Group, Day, and all interaction terms, and included subject as a random intercept to account for repeated-measures. We first explored potential changes in calcium activity during correct choices (**Figure 3c**). Our AUC model revealed no significant fixed effects of Epoch (F(1,154.82)=0.06, p=0.8), Group (F(2,136.92)=0.03, p=0.98), or Day (F(1,156.94)=0.78, p=0.38) or significant interactions, including Epoch*Group (F(2,154.82)=0.07, p=0.94), Epoch*Day (F(1,155.30) = 0.58, p=0.45), Group*Day (F(2,156.84)=0.78, p=0.46), Epoch*Group*Day (F(2,155.29)=0.66, p=0.52). These findings indicate that potential changes in calcium activity were not driven by Epoch, Group, or by Day. To further explore learning-related changes in calcium, we estimated AUC slopes across acquisition and found no significant changes in calcium signal for Control (β=0.108 ± 0.073, 95% CI [-0.037, 0.253]), Hapln1 (β=−0.006 ± 0.071, 95% CI [-0.146, 0.133]) or Gi (β=0.004 ± 0.065, 95% CI [-0.125, 0.133]) groups.

Next we explored incorrect responses (**Figure 3d**). Our model revealed no significant fixed effects for Epoch (F(1,158.05)=0.09, p=0.77), Group (F(2,22.69)=0.351, p=0.71), or Day (F(1,158.20)=0.47, p=0.49) and no significant interactions, including Epoch*Group (F(2,158.05)=0.08, p=0.92), Epoch*Day (F(1,158.07)=0.002, p=0.97), Group*Day (F(2,158.20)=0.43, p=0.65), or Epoch*Group*Day (F(2,158.07)=0.001, p=0.99). Slope analysis revealed that each groups calcium activity was stable during incorrect choices and did not differ across days: Control (β=0.036 ± 0.369, 95% CI [-0.692, 0.764]), Gi (β=0.389 ± 0.323, 95% CI [-0.249, 1.027]), and Hapln1 (β=−0.011 ± 0.350, 95% CI [-0.703, 0.681]). Lastly, we compared the difference between AUCs from the last day to the first day of learning within groups and between groups for both correct and incorrect choices. We found no differences for both correct and incorrect choices within Controls (p=0.38; p=0.93), Hapln1 (p=0.86; p=75), or Gi (p=0.86; p=0.23) or between group differences (p=0.518, ANOVA; p=0.65, ANOVA) for changes in AUCs (**Supplementary Figure 2**). Together, these results demonstrate that DREADD and Hapln1 manipulation did not influence acquisition performance or motivation, as animals showed no differences in time to access the reward or alter calcium activity during both correct and incorrect responses. Lastly, LCN PV cell calcium activity remained consistent across learning and groups.

### DREADD and Hapln1 Manipulation Alters Reversal Learning

To analyze reversal performance, we conducted the same logit transformation as acquisition and first compared all logits using a Two-way ANOVA with factors Group and Day. Our results revealed no Group differences in logits across all reversal days (F(2,91)=1.78, p=0.18, ANOVA), however, there was a main effect of Day (F(1,91)=14.89, p=0.0002, ANOVA). Slope logits analysis revealed group differences (F(2,16)=5.64, p=0.014, ANOVA), with Gi animals showing an increase in reversal slope compared to Controls (p=0.01, Tukey). There were no slope differences between Gi and Hapln1 (p=0.38, Tukey) or between Controls and Hapln1 (p=0.17, Tukey) groups. In analyzing the last day of reversal, group comparisons (F(2,16)=3.95, p=0.04, ANOVA) revealed Gi animals showed in increase in performance compared to Control (p=0.002, Tukey) and Hapln1 animals (p=0.036, Tukey) (**Figure 4a**). Our latency analysis revealed no group differences in trial initiation latency (F(2,16)=2.1, p=0.16, ANOVA), responding to the stimulus latency (F(2,16)=0.69, p=0.51, ANOVA), or collecting a reward latency (F(2,16)=1.4, p=0.27, ANOVA) (**Supplementary Figure 3)**. Next, we explored whether the number of completed trials was related to reversal performance and found that trials were not a significant predictor (p=0.55) of performance. Further exploration of trial completion revealed group differences in the cumulative sum (F(2,16)=3.67, p=0.049, ANOVA), specifically Hapln1 animals completing more trials than Controls (p=0.039, Tukey) with no differences in Gi (p=0.34, Tukey) or between Gi and Controls (p=0.38, Tukey) (**Figure 4b**). Lastly, we analyzed the relationship between the performance on the last day of reversal and DREADD expression across the CN. We found that CN DREADD expression was not correlated (R=0.45, p=0.26) (**Supplementary Figure 4)**.

**Figure 4.**
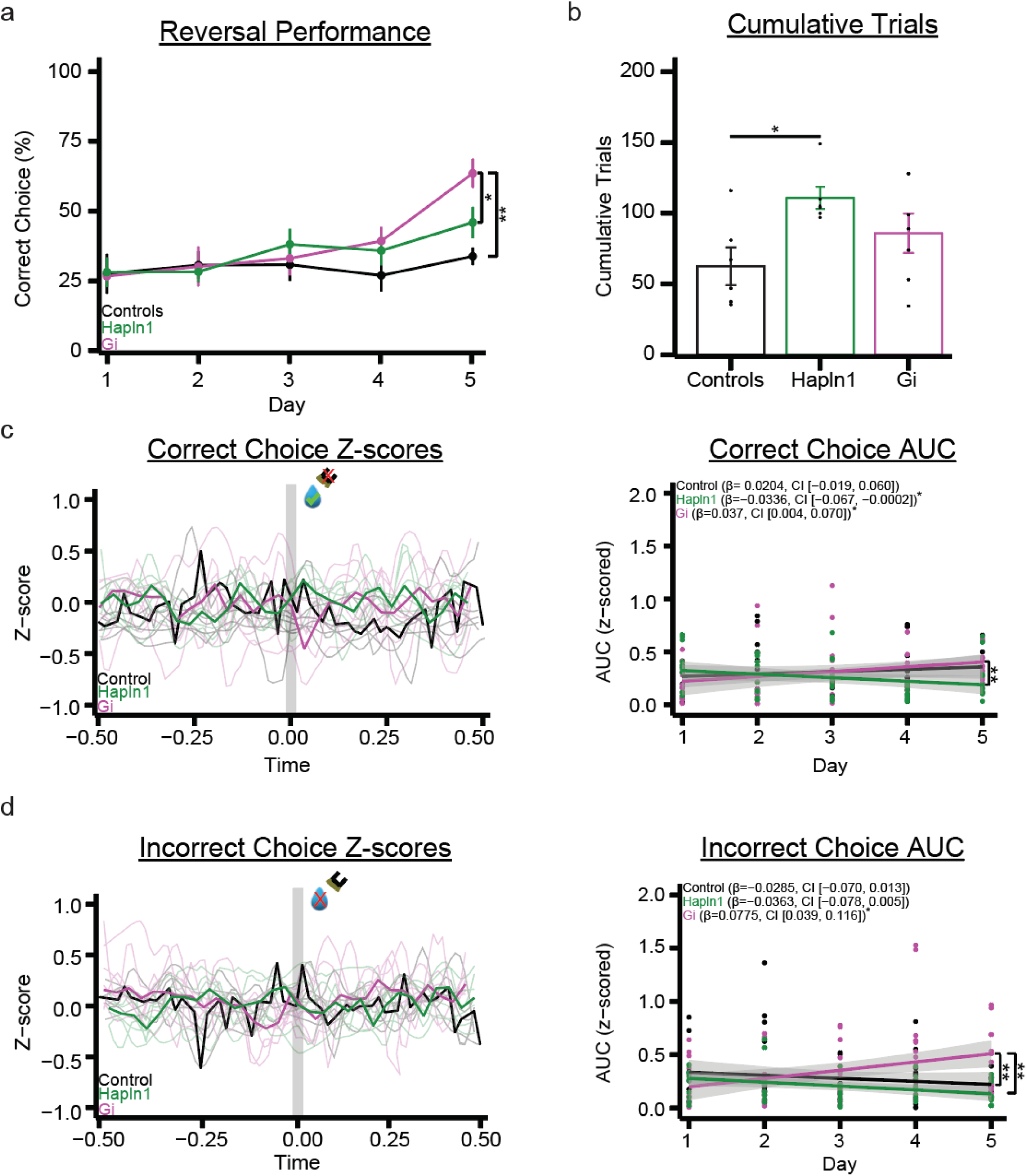
Reversal learning is impacted by both inhibitory DREADDs and Hapln1. Control (black, n=6), Hapln1 (green, n=6), Gi (magenta, n=7). (A.) Line graphs of correct choice (%) during the visual discrimination reversal stage in male mice. A logit transformation of reversal correct choice (%) was performed and a linear regression using the logit scores was then performed to calculate the slopes and analyzed using Two-way ANOVAs with group and day as a factor. Mice showed no group differences in all logits (F(2, 91)=1.78, p=0.18, ANOVA) or slopes (F(2,16)=0.8, p=0.47, ANOVA). The analysis of the final day reversal logits revealed group differences (F(2,16)=3.949, p=0.04), driven by Gi animals increased performance compared to controls (p=0.002, Tukey) and Hapln1 (p=0.036, Tukey) groups. (B.) Comparison of group cumulative completed trials across reversal (F(2,16)=3.678, p=0.048) revealed Hapln1 animals completed more trials overall compared to controls (p=0.039, Tukey). (C.) (Left) Visualized GCaMP z-scores from 0.5 s before and 0.5 s after a correct choice occurred. Vertical grey bar indicates at timepoint zero when a correct response was made and semitransparent lines are averaged correct choice z-scores per animal with bold lines indicating group averages. (Right) Area-under-the-curve (AUC) slope analysis for correct choice was the same as acquisition. Simple Slope analysis revealed group-dependent changes in calcium activity, specifically an increase in correct-choice related calcium activity in Gi animals (β=0.037, CI [0.004, 0.070]) and a decrease in correct-choice related calcium activity in Hapln1 animals (β=−0.0336, CI [−0.067, −0.0002]). Control animals showed no differences in correct choice related calcium activity (β= 0.0204, CI [−0.019, 0.060]). Group slope comparison revealed differences between Gi and Hapln1 (p=0.001, Tukey) animals, and no differences between Gi and Control (p=0.8, Tukey) or Control and Hapln1 (p=0.1, Tukey) animals. (D.) (Left) Visualized GCaMP z-scores from 0.5 s before and 0.5 s after an incorrect choice was performed. Vertical grey bar indicates at timepoint zero when an incorrect choice occurred and semitransparent lines are averaged incorrect choice z-scores per animal with bold lines indicating group averages. Simple Slope analysis revealed group-dependent changes in calcium activity, specifically an increase in incorrect-choice related calcium activity in Gi animals (β=0.0775, CI [0.039, 0.116]). Data represents mean ± SEM, *p < 0.05, **p < 0.01

Together, these findings indicate that reversal performance improved across days similarly across groups, but DREADD animals exhibited improved performance specifically on the final day of reversal. These differences could not be explained by latency measures or trial completion, but rather Gi manipulation may facilitate reversal learning specifically. Moreover, Hapln1 treated animals showed a slight improvement in reversal learning, albeit not statistically significant, compared to controls.

### Group-dependent Changes in Reversal Correct Choice Calcium Activity

To analyze reversal calcium activity, we conducted the same analysis as with acquisition using LLMs with fixed effects of Epoch, Group, Day, and all interaction terms, and subject as a random effect for both correct and incorrect choice instances for reversal. Starting with correct choice, AUC analysis revealed no significant main effects for Epoch (F(1,144)=0.08, p=0.53), Group (F(2,54)=1.45, p=0.24)), or Day (F(1,149)=0.58, p=0.25) and no significant interactions involving Epoch, such as Epoch*Group (F(2,148.44)=1.1, p=0.34), Epoch*Day (F(1,144.18)=0.08, p=0.77), or Epoch*Group*Day significant (F(2,144)=0.88, p=0.42). However, a significant Group*Day interaction (F(2,148.4) = 4.74, p=0.01) was detected, indicating that changes in calcium signal across days differed as a function of Group (**Figure 4c**).

To further characterize the significant Group*Day interaction, we performed simple-slopes analyses using estimated marginal trends to examine the effect of Day within each group. This analysis revealed Gi mice exhibited a significant positive change in AUC across days (β=0.037, 95% CI [0.004, 0.070]), Hapln1 mice exhibited a significant negative change in AUC across days (β=−0.0336, 95% CI [−0.067, −0.0002]), and Control mice showed no significant change in AUC across days (β=0.0204, 95% CI [−0.019, 0.060]). Pairwise comparisons of slopes confirmed that the Gi animals differed significantly from the Hapln1 group (p=0.01, Tukey), whereas neither group differed significantly from controls (Gi, p=0.8, Tukey; Hapln1, p=0.1, Tukey) (**Figure 4c**).

Together, these results indicate that correct choice-related changes in calcium activity are group-dependent, with Gi animals showing a progressive increase in calcium signal, Hapln1 animals showing a progressive decrease, and controls remaining stable over time. Importantly, these effects were independent of Epoch, suggesting that group differences primarily reflect longer-timescale adaptations rather than acute pre-versus post-event mechanisms.

### DREADD Manipulation Increase Reversal Incorrect-Choice Calcium Activity

Next, we performed LLMs for incorrect choice instances during reversal. Our model revealed no significant fixed effects for Epoch (F(1,162)=0.27, p=0.6), Group (F(2,76.42)=2.65, p=0.07), or Day (F(1,162)=0.13, p=0.72) or interactions, including Epoch*Group (2,162)=0.03, p=0.97), Epoch*Day (F(1, 162)=0.48, p=0.49), and Epoch*Group*Day (F(2,162)=0.28, p=0.75). We did find a significant Group*Day interaction (F(3,159)=7.15, p=0.0001), indicating that changes in calcium signal across training days differed significantly between groups. At the level of fixed-effect coefficients, this interaction was driven by a significant positive Day-related change in the Gi group (p=0.002), whereas no significant Day-related effects were observed for the Control (p=0.87) or Hapln1 (p=0.98) groups. To further characterize the significant Group*Day interaction, simple-splopes analysis revealed that Gi mice exhibited a significant positive increase in AUC across days (β=0.0775, 95% CI [0.039, 0.116]), Control mice showed no significant change in AUC across days (β=−0.0285, 95% CI [−0.070, 0.013]), and Hapln1 mice also showed no significant change in AUC across days (β=−0.0363, 95% CI [−0.078, 0.005]). Pairwise comparisons of slopes confirmed that the Gi animals differed significantly from both Control (p=0.001, Tukey) and Hapln1 mice (p=0.0003, Tukey), whereas Control and Hapln1 groups did not differ from one another (p=0.96, Tukey) (**Figure 4d**). Lastly, we compared the difference between AUCs value from the last day to the first day of learning within groups and between groups for both correct and incorrect choices. We found no differences for both correct and incorrect choices within Controls (p=0.38; p=0.93), Hapln1 (p=0.86; p=75), or Gi (p=0.86; p=0.23) groups or between group differences (p=0.518, ANOVA; p=0.65, ANOVA) for changes in AUCs (**Supplementary Figure 5)**. Finally, we performed a multiple regression model predicting reversal logit slopes using correct choice and incorrect AUC slopes to explore the potential relationship between reversal learning and PV cell activity. Our model revealed that correct choice AUC slopes (p=0.46) and incorrect choice AUC slopes (p=0.91) were not significant predictors for reversal performance (**Figure 4d**).

Overall, these results demonstrate a progressive increase in calcium activity during incorrect choices in Gi animals compared to stable activity observed in Control and Hapln1 animals. Additionally, this group specific change was not reflected in last to first day AUC differences. These findings suggest that Gi manipulation specifically altered error-related neural signaling across learning compared to Control and Hapln1 animals, but the altered signal alone was not sufficient to explain the changes in reversal learning.

### Adolescent LCN Manipulation Alters PNN Link Protein Hapln1

After animals completed reversal between P65 and P70, we collected cerebellar tissue to stain and analyze PNN link protein, Hapln1 within the LCN target region across groups (Figure 5a, **Supplementary Table 1**). We first compared LCN Hapln1 cell counts across groups and found no differences in the amount of Hapln1 expressing cells normalized by area (F(2,62)=1.08, p=0.35, ANOVA) (**Figure 5b**). We then performed a linear regression to predict LCN Hapln1 intensity using Group as a predictor (F(2,62)=5.76, p=0.005; R^2^=0.13). Our model revealed Gi membership is estimated to increase LCN Hapln1 intensity by 13% (p=0.01) and Hapln1 vector manipulation is estimated 20% (p=0.003). (**Figure 5c**). Our post hoc comparisons revealed Hapln1 manipulated animals (n=6) had increased Hapln1 expression compared to Controls (p=0.007, Tukey), as well as a slight increase in Gi animals (n=6) Hapln1 expression compared to Controls (p=0.03, Tukey). There were no differences between Hapln1 and Gi groups (p=0.9, Tukey). Finally, we performed a Pearson correlation to identify any potential relationships between LCN Hapln1 and reversal performance. Our results revealed no relationship between Hapln1 and final day reversal performance (R²=0.18 p=0.08) (**Figure 5d**).

**Figure 5.**
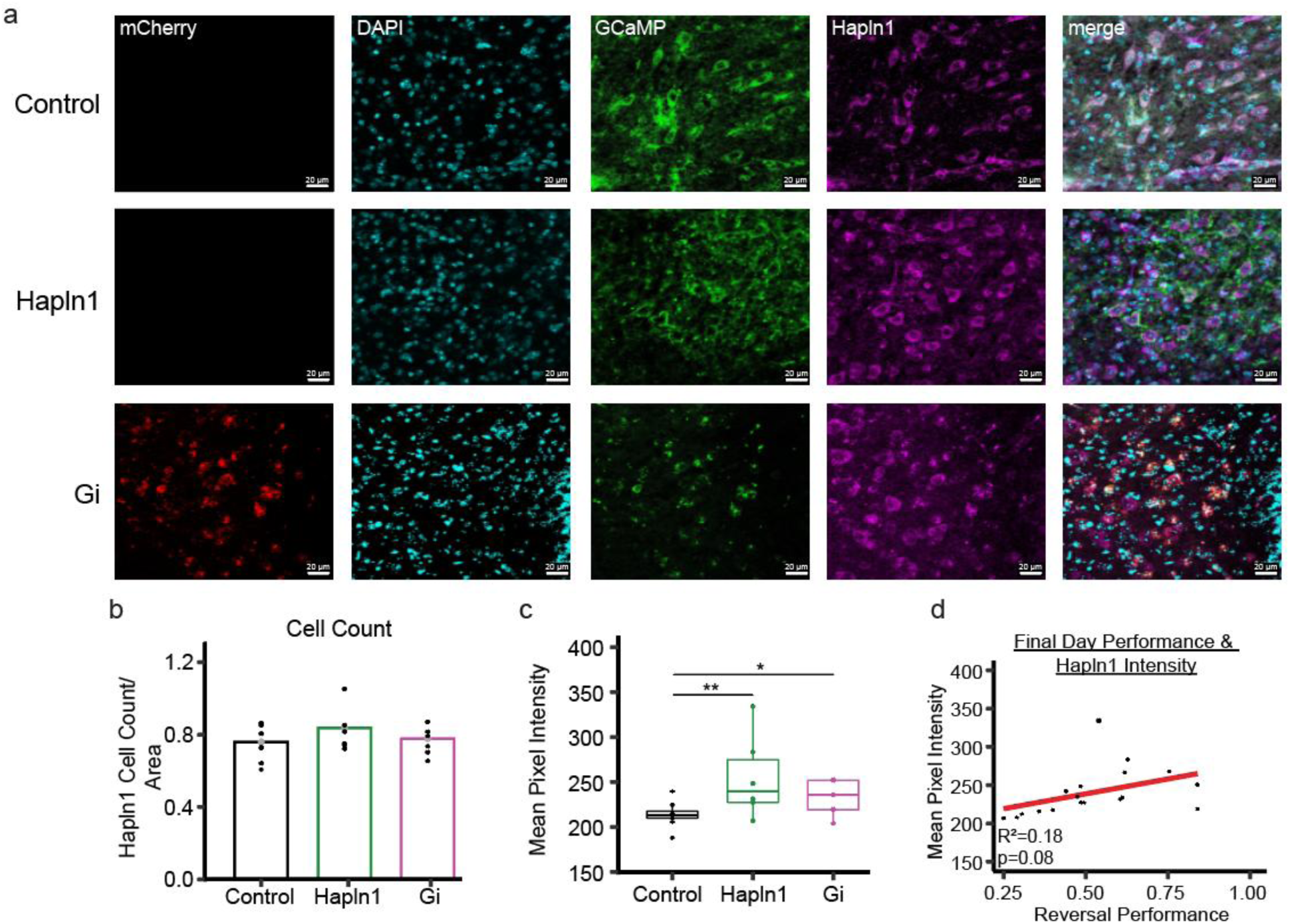
Enhancement of PNN link protein Hapln1. (A.) Immunofluorescence shows Gi DREADDs (mCherry), GCaMP (green), and extracellular Hapln1 positive cells stained for anti-Hapln1 (magenta) in the lateral cerebellar nucleus (LCN) in Control, Hapln1, and Gi animals (Scale bar: 20 μm, taken at 40x objective) (B.) As expected, there were no differences in Hapln1 cell count between groups (F(2,62)=1.08, p=0.35, ANOVA). (C.) LCN Hapln1 intensity differed significantly between groups (F(2,62)=5.76, p=0.005, ANOVA). Post hoc analysis revealed Hapln1 overexpressing animals (p=0.007, Tukey) had increased anti-Hapln1 intensity compared to Controls. Also, Gi animals (p=0.03, Tukey) showed increased Hapln1 intensity compared to Controls. (D.) Pearson correlation between Hapln1 intensity and reversal performance. There was no significant relationship identified between Hapln1 and reversal performance (R²=-0.18 p=0.08). Data represents mean ± SEM, *p < 0.05, **p < 0.01

Together, these findings suggest that Hapln1 vector and DREADD manipulation did not alter Hapln1 expressing cell quantity, but rather influenced the Hapln1 intensities compared to controls. Additionally, there was no relationship identified between reversal performance and LCN Hapln1 intensities, suggesting that subtle changes in LCN Hapln1 expression may not be sufficient to influence the dynamic processes that underlie cognitive behavior.

### LCN Manipulations on Aggrecan

To further explore LCN PNN components and their relationship with behavior, we analyzed LCN aggrecan expression across groups (**Supplementary Table 2**), as aggrecan is the most abundant CPGS across PNN populations ^35^ (**Supplementary Figure 6a**). First, we compared LCN aggrecan positive cell counts across groups. We found no significant group differences (F(2,35)=1.09, p=0.35, ANOVA), suggesting DREADD and Hapln1 manipulations did not alter aggrecan expressing cell counts (**Supplementary Figure 6b**). We then performed a linear regression to predict LCN aggrecan intensity using Group as a predictor variable and found our overall model was significant (F(2,35)=3.76, p=0.03, R²=0.18), indicating that aggrecan intensity varied as a function of Group. However, comparisons to the Control group were not significant, with Gi animals showing a small decrease by 18% (p=0.14) and Hapln1 animals showing a slight increase by 12% (p=0.34). As our model indicated group differences, we performed post hoc comparisons and found Hapln1 animals showed an increase in aggrecan intensity compared to Gi animals (p=0.03, Tukey) (**Supplementary Figure 6c**). Lastly, we performed a Pearson correlation to explore whether aggrecan intensity within the LCN was related to reversal performance. Our results revealed that aggrecan intensity was unrelated to reversal performance (R²=-0.092 p=0.68) (**Supplementary Figure 6d)**.

Overall, Hapln1 and Gi manipulation did not alter aggrecan cell counts or aggrecan intensities, and LCN aggrecan intensity was unrelated to reversal performance. There was however, a difference between Gi and Hapln1 aggrecan intensities, indicating the DREADD manipulation could be affecting alternative PNN related mechanisms that influence CSPGs.

## Discussion

In the current study, we investigated the role of the LCN on cognitive behavior during a critical period of adolescent development. The use of chemogenetic inhibition of LCN neuronal populations and a viral vector approach to increase PNN link protein Hapln1, yielded impacts on learning and neural activity of PV+ cells. Gi DREADDs enhanced reversal learning and there was increased calcium activity during incorrect trials, suggesting a type of teaching signal response that benefits performance in this task. We have previously shown that Gi DREADD mice have reduced PNNs after reversal learning, suggesting an increase in brain plasticity that benefits learning (Lyle et al., 2025). On the other hand, Hapln1 mice required more trials compared to controls in both acquisition and reversal, and had less calcium responses after choosing a correct and incorrect choice in reversal compared to Gi mice. These findings suggest that ECM remodeling in the LCN can impact learning of the visual discrimination task.

While all animals were able to learn the visual discrimination task and there were no differences in calcium activity during acquisition, reversing presented a greater challenge. Reversal learning revealed group-specific effects on adolescent LCN manipulation. Gi animals demonstrated enhanced reversal learning, evident by increased learning slopes and improved performance on the final reversal of reversal, whereas Hapln1 overexpressing animals showed a slight increase in performance, albeit insignificant. Of importance, the differences in performance were not explained by trial completion or latency measures, indicating that inhibitory LCN DREADD manipulation during adolescence selectively facilitated reversal learning, consistent with previous research.^2^ Analysis of reversal calcium recordings revealed a unique relationship with learning. Calcium activity during correct choices was differentially modulated across groups with Gi animals displaying a progressive increase in correct choice dependent activity whereas Hapln1 animals showed a decrease compared to Controls. Moreover, Gi manipulation also increased incorrect choice calcium responses, demonstrating that adolescent LCN inhibition may amplify error-related activity, thus aiding in reversal learning. Interestingly, these changes were independent of pre- and post-response events, suggesting that changes in calcium activity reflect longer timescale plasticity and which may have also impacted cerebello-cortical circuits. ^10,11^

Previous work in our lab has found that adolescent LCN inhibition in male mice improves reversal learning and decreases global LCN PNN intensities stained with Wisteria floribunda agglutinin (WFA).^2^ Therefore, we further explored PNN mechanism influence on reversal learning by analyzing indispensable PNN components, link protein Hapln1 and one component of the PNNs, aggrecan.^32,36,45^ It is important to note that aggrecan binds to hyaluronan through a N-terminal G1 domain and Hapln1 stabilizes this interaction, but Hapln1 does not directly influence the amount of aggrecan present^46^. Analysis of aggrecan revealed no changes to aggrecan intensities or cell counts as a result of Hapln1 overexpression. Gi Gi mice had reduced intensity of aggrecan compared to Hapln1 mice, but no change compared to controls. This is interesting as Gi mice have reduced WFA, therefore more studies are necessary to determine changes in PNN composition. These results indicate that DREADD and Hapln1 manipulations resulted in remodeling of the ECM.

Our findings suggest that adolescent LCN inhibition improves behavioral flexibility and neural plasticity during reversal learning. Specifically, LCN inhibition may facilitate cognitive adaptations by increasing correct and error-related PV cell activity. Whereas Hapln1 overexpression may close critical periods earlier by increasing the probability of CSPG and functionality of CSPGs, thus constraining flexibility and limiting neural plasticity during reversal learning. As LCN Hapln1 and aggrecan cell expression were not correlated to reversal learning, we cannot rule out differences in synthesis rates or proteolysis, therefore future research will explore alternative LCN PNN components and other mechanisms following critical period perturbations. For example, brevican is a part of the CSPG family of lecticans, and is found to limit the mobility of potassium channels and AMPARs in PV-expressing interneurons. Moreover, deletions in brevican enhance PV cell excitability by decreasing the action potential threshold and latency to fire. Additionally, others^47^ have found that brevican increases around PV cells into the third postnatal week in the hippocampus, therefore, it is possible that LCN neuronal inhibition at P21 impacted brevican expression, which could explain the increased calcium response during reversal in Gi animals. Other PNN mechanisms, such as Tenascin-R, a PNN component that crosslinks lecticans and contributes to the structural integrity and rigidity of the net,^48^ or the sulfation pattern of chondroitin sulfate glycosaminoglycan chains which critically influences PNN function by modulating interactions with growth factors, guidance cues, and synaptic adhesion molecules ^49^ can also be explored in relation to LCN plasticity and learning. Finally, within the mouse LCN, inhibitory interneurons make up approximately 11% of the neuronal population,^50^ so future studies can analyze excitatory populations as well. However, our findings did identify increases in PV activity across reversal, which suggest that PV cells in Gi animals had increased inhibition onto LCN pyramidal output neurons, thereby adjusting cerebellar error-signals to the cerebello-cortical circuit to enhance reversal learning. Future research should record changes in pyramidal cell activity following LCN perturbations during reward and related responses and measure cortical activity to explore potential connectivity changes in the cerebello-cortico circuit.

Together, these findings suggest that adolescent LCN inhibition signaling promotes sustained amplification of task-related PV calcium activity across reversal learning, whereas Hapln1 manipulation may precociously dampen this activity through ECM remodeling, thereby reflecting differential impacts on LCN plasticity or PNN-dependent limiting factors on PV cell activity.

## Acknowledgements

This work was supported by: the Institute for Mental Health Research, Institute for Social Science Research, Nancy Eisenberg Junior Faculty scholar Award, Arizona Department of Health Sciences (ADHS14-052688), US Department of Health and Human Services and the state of Arizona (ADHS Grant No. CTR057001), National Institute on Aging (NIA) of the National Institutes of Health (NIH) (P30AG019610), Arizona Alzheimer’s Disease Research Center REC Fellows Program and Arizona Alzheimer’s Consortium. Figures were created with Biorender.com.

## Author Contributions

T.L.: Conceptualization, data analysis and curation, validation, investigation, visualization, methodology, writing, and project administration. A.B.: Data analysis. J.L.V.: Conceptualization, data analysis and curation, funding acquisition, validation, investigation, visualization, methodology, writing, and project administration.

## Data and Code Availability

Lyle, Tristan; Berkley, Aaron; Verpeut, Jessica, 2026, “Replication Data for: Elucidating the role of cerebellar nuclei parvalbumin activity on adolescent reversal learning”, https://doi.org/10.48349/ASU/KKWFWE, ASU Library Research Data Repository, V1

## Code availability

All experimental and analysis code is available at Arizona State University dataverse.

## Supplemental information

Table S1-2.

Figures S1–6

Data S1. Raw experimental data containing behavior, immunohistochemistry, and calcium imaging for each experiment group (CNO-only, Gi, Hapln1).

Data S2. Statistical comparisons between groups control (CNO-only), DREADD condition (Gi), and Hapln1 for behavior, immunohistochemistry, and calcium imaging data.

## Supplementary

**Supplementary Table 1.**
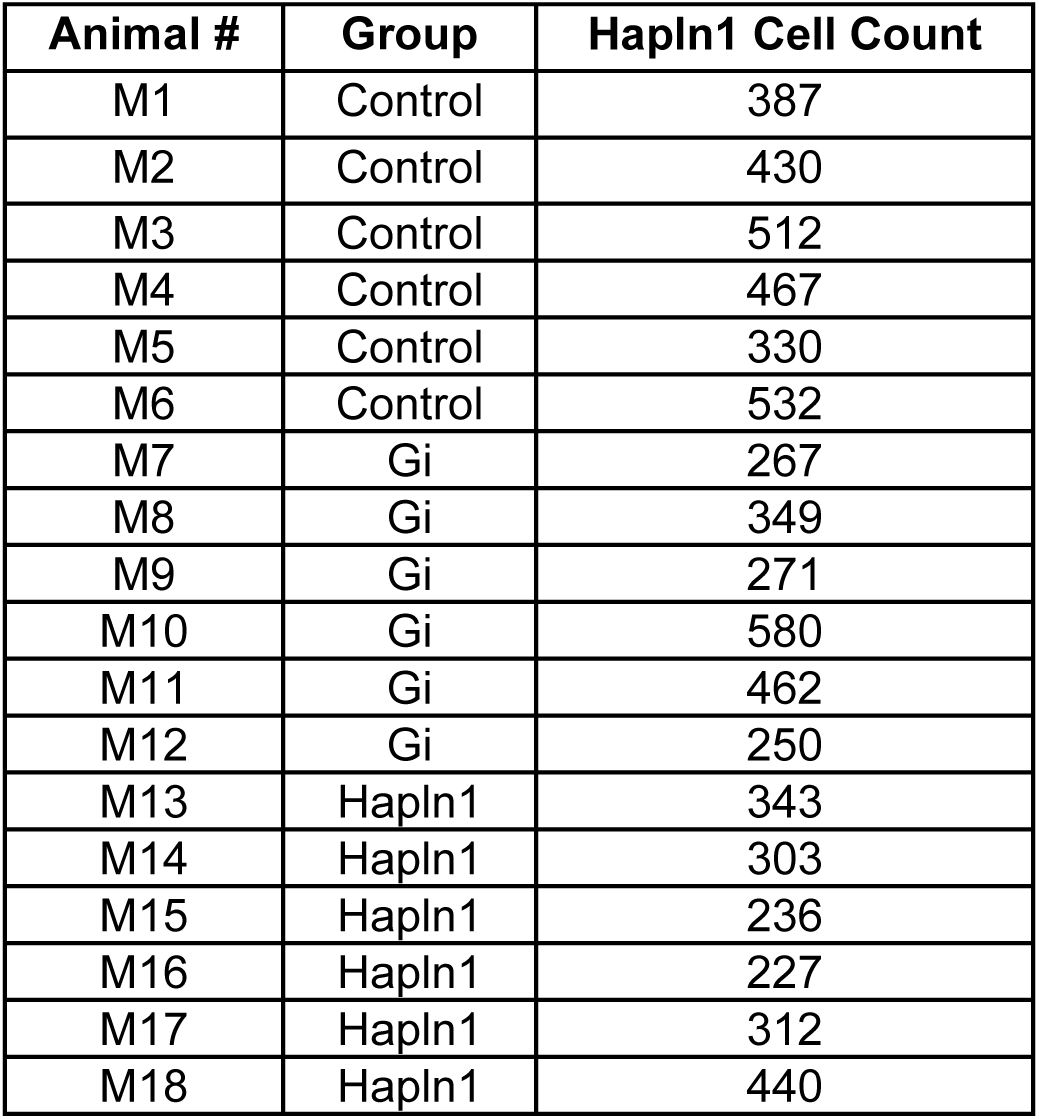
Hapn1 cell count by mouse and group (n=6 per group).

| Animal # | Group | Hapln1 Cell Count |
| --- | --- | --- |
| M1 | Control | 387 |
| M2 | Control | 430 |
| M3 | Control | 512 |
| M4 | Control | 467 |
| M5 | Control | 330 |
| M6 | Control | 532 |
| M7 | Gi | 267 |
| M8 | Gi | 349 |
| M9 | Gi | 271 |
| M10 | Gi | 580 |
| M11 | Gi | 462 |
| M12 | Gi | 250 |
| M13 | Hapln1 | 343 |
| M14 | Hapln1 | 303 |
| M15 | Hapln1 | 236 |
| M16 | Hapln1 | 227 |
| M17 | Hapln1 | 312 |
| M18 | Hapln1 | 440 |

**Supplementary Table 2.**
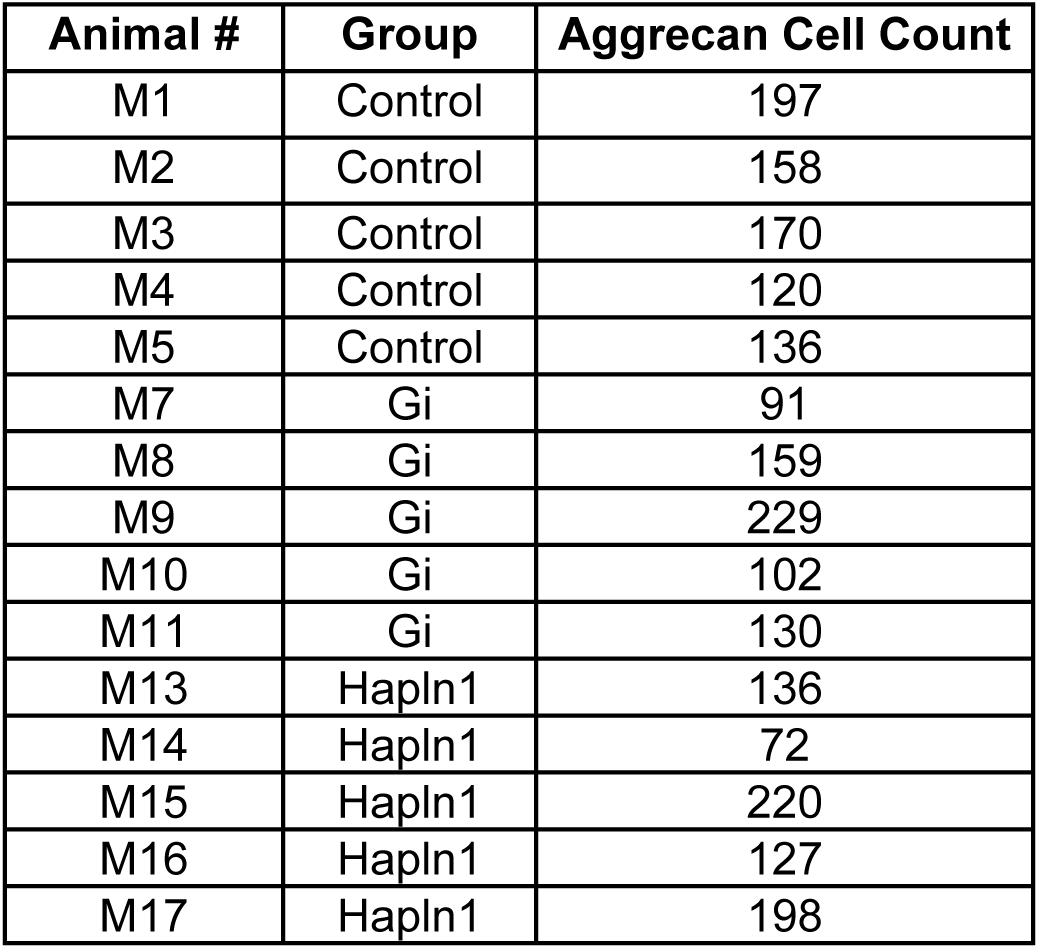
Aggrecan cell count by mouse and group (n=5 per group).

| Animal # | Group | AggreCAN Cell Count |
| --- | --- | --- |
| M1 | Control | 197 |
| M2 | Control | 158 |
| M3 | Control | 170 |
| M4 | Control | 120 |
| M5 | Control | 136 |
| M7 | Gi | 91 |
| M8 | Gi | 159 |
| M9 | Gi | 229 |
| M10 | Gi | 102 |
| M11 | Gi | 130 |
| M13 | Hapln1 | 136 |
| M14 | Hapln1 | 72 |
| M15 | Hapln1 | 220 |
| M16 | Hapln1 | 127 |
| M17 | Hapln1 | 198 |

**Supplementary Figure 1.**
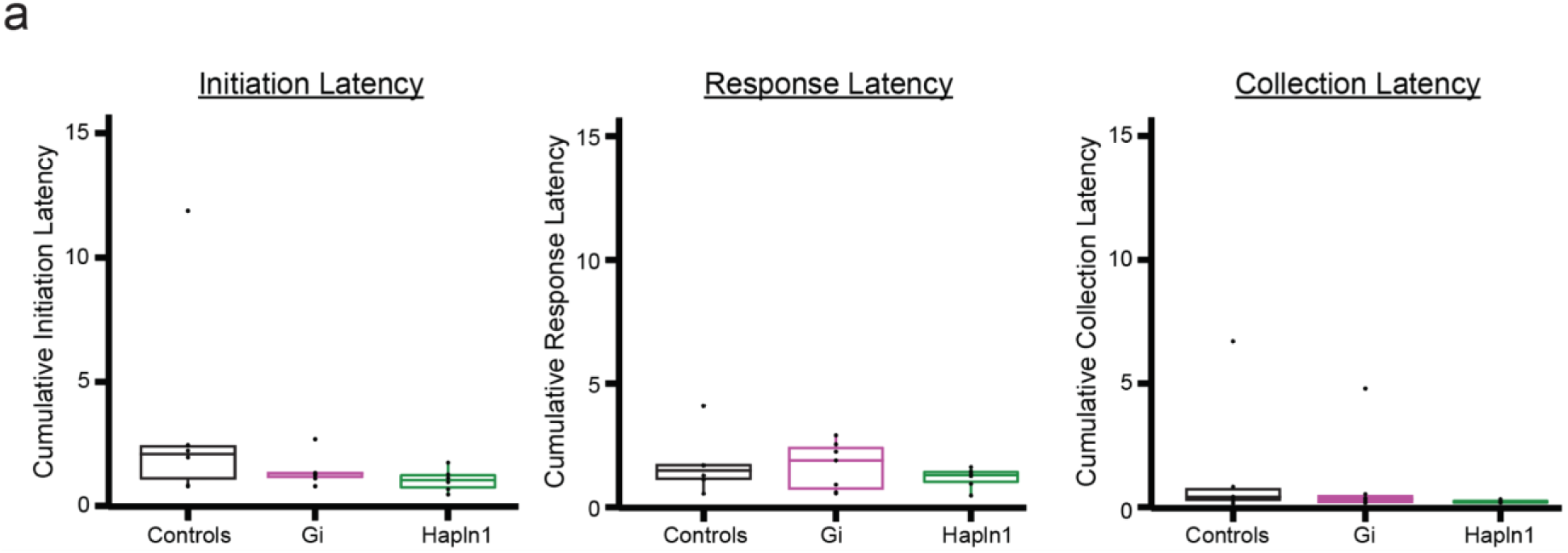
Acquisition latencies. (A.) No group differences were found in initiation (F(2,16)=1.63, p=0.23, ANOVA), response (F(2,16)=0.64, p=0.54, ANOVA), or collection latencies ((F(2,16)=0.73, p=0.49, ANOVA) in seconds. Data represents mean ± SEM.

**Supplementary Figure 2.**
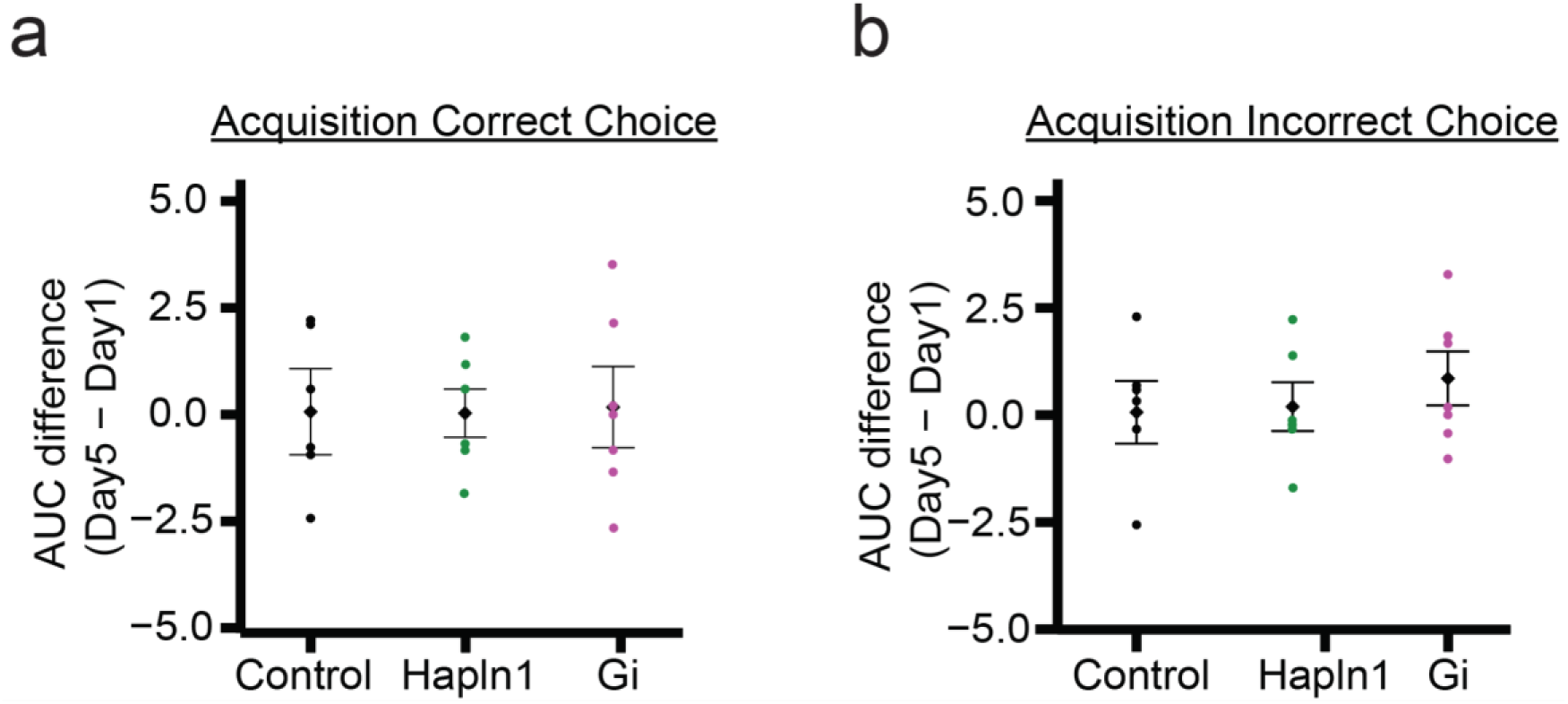
Acquisition calcium activity. (A.) Difference plot between day 5 and day 1 AUC values per group (control, black; Hapln1, green; Gi, magenta) for correct choices. There are no significant differences between groups AUC difference scores (F(2,16)=0.69, p=0.52, ANOVA). (B.) Incorrect choice difference plot between day 5 and day 1 AUC values per group with no significant differences found (F(2,16)=0.45, p=0.65, ANOVA). Data represents mean ± SEM.

**Supplementary Figure 3.**
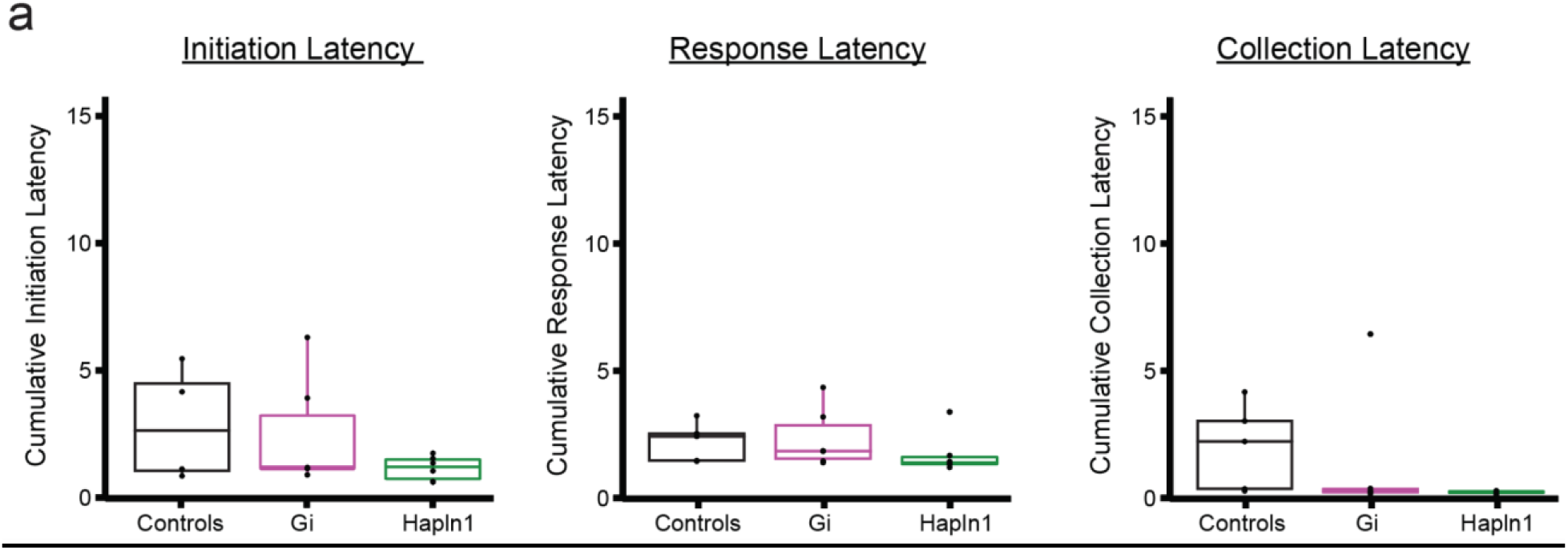
Reversal latencies. (A.) No group differences were found in animals initiation (F(2,16)=2.1, p=0.16, ANOVA), response (F(2,16)=0.7, p=0.51, ANOVA), or collection latencies (F(2,16)=0.73, p=0.27, ANOVA). Data represents mean ± SEM.

**Supplementary Figure 4.**
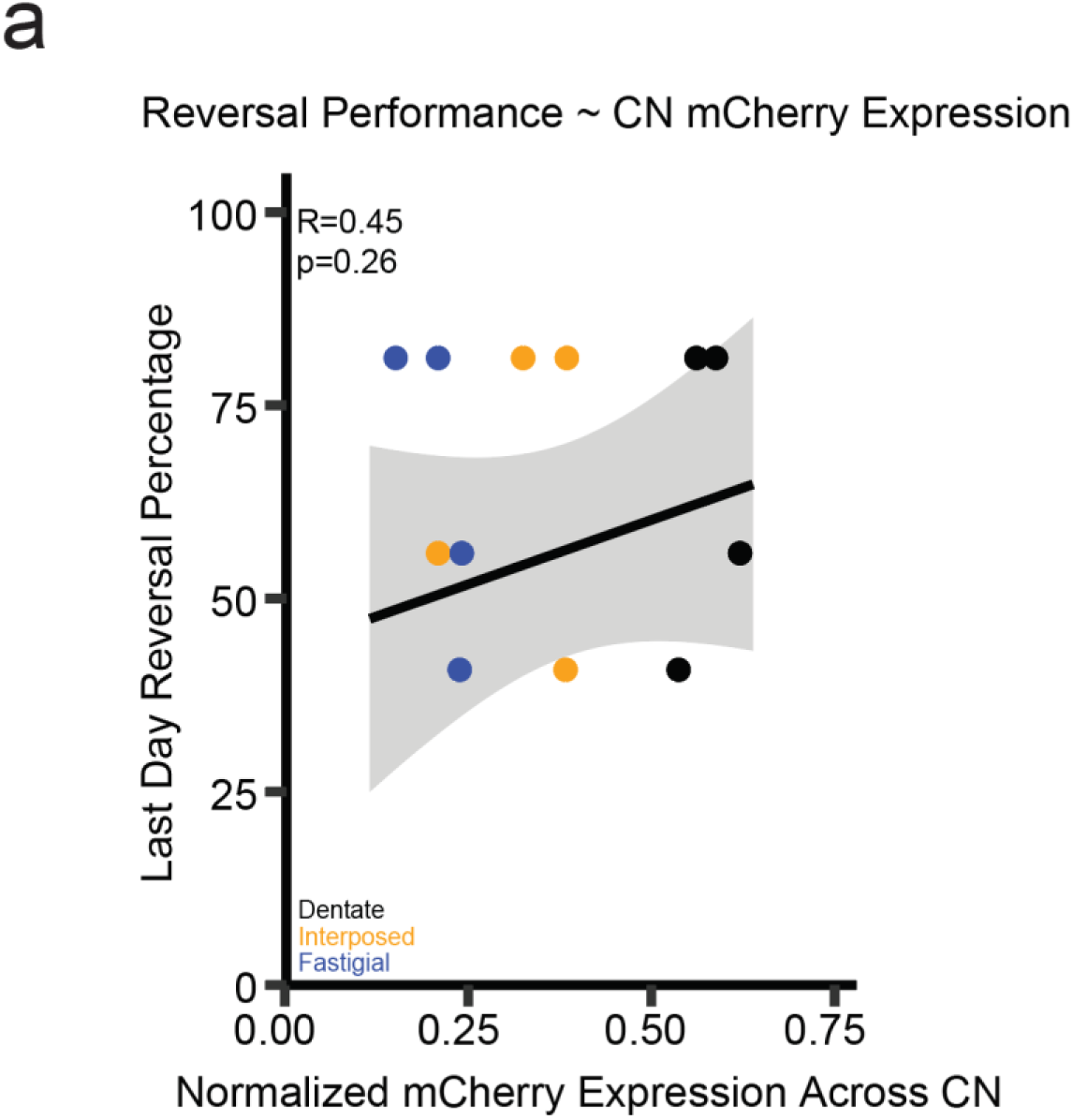
DREADD correlations to reversal performance. (A.) Scatterplot of DREADD reporter expression within the CN (dentate, black; interposed, orange, fastigial, blue). and the last day of reversal performance. Pearson correlation revealed no significant relationship (p=0.26, R=0.2, Pearson Correlation) (n=4, Gi).

**Supplementary Figure 5.**
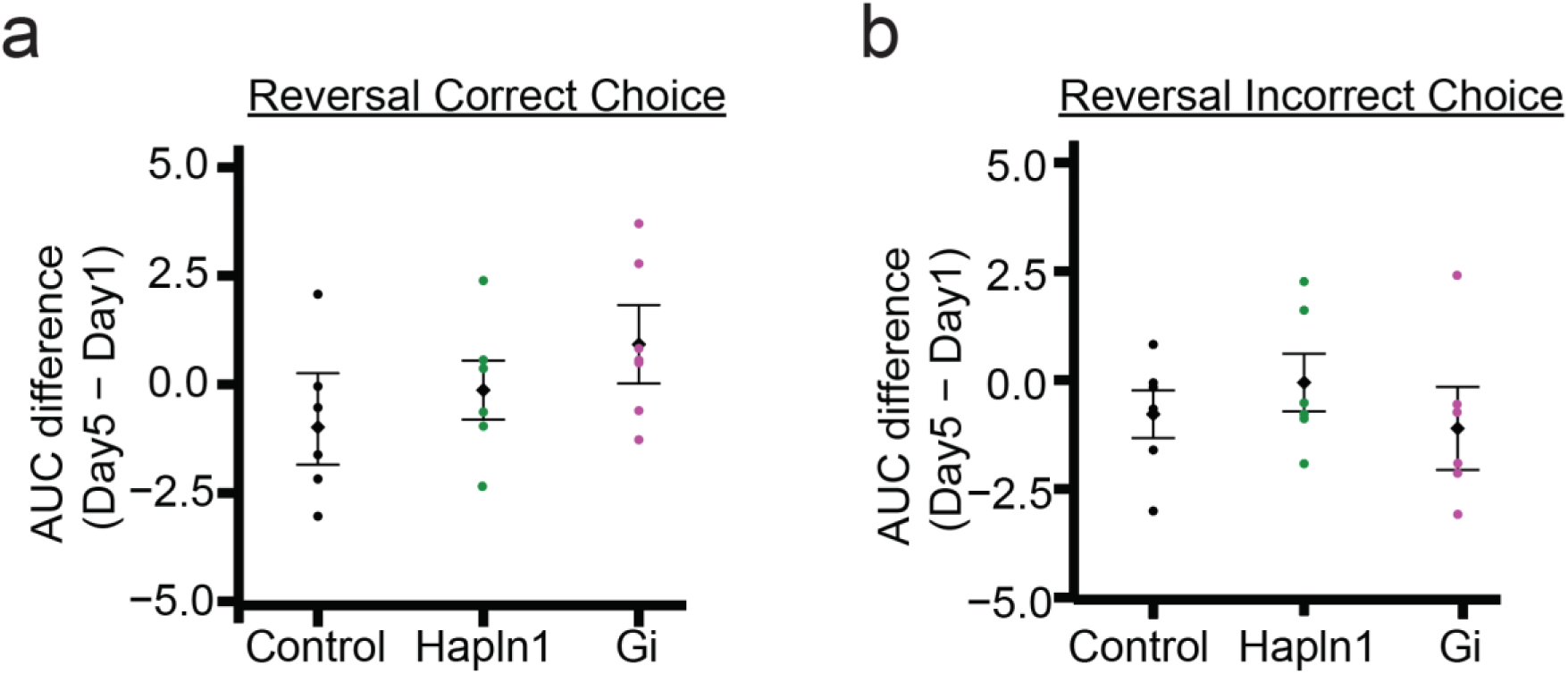
Reversal calcium activity. (A.) Difference plot between day 5 and day 1 AUC values per group (control, black; Hapln1, green; Gi, magenta) for correct choices. There are no significant differences between groups AUC difference scores (F(2,16)=0.81, p=0.46, ANOVA). (B.) Incorrect choice difference plot between day 5 and day 1 AUC values per group with no significant differences found (F(2,16)=0.55, p=0.59, ANOVA). Data represents mean ± SEM

**Supplementary Figure 6.**
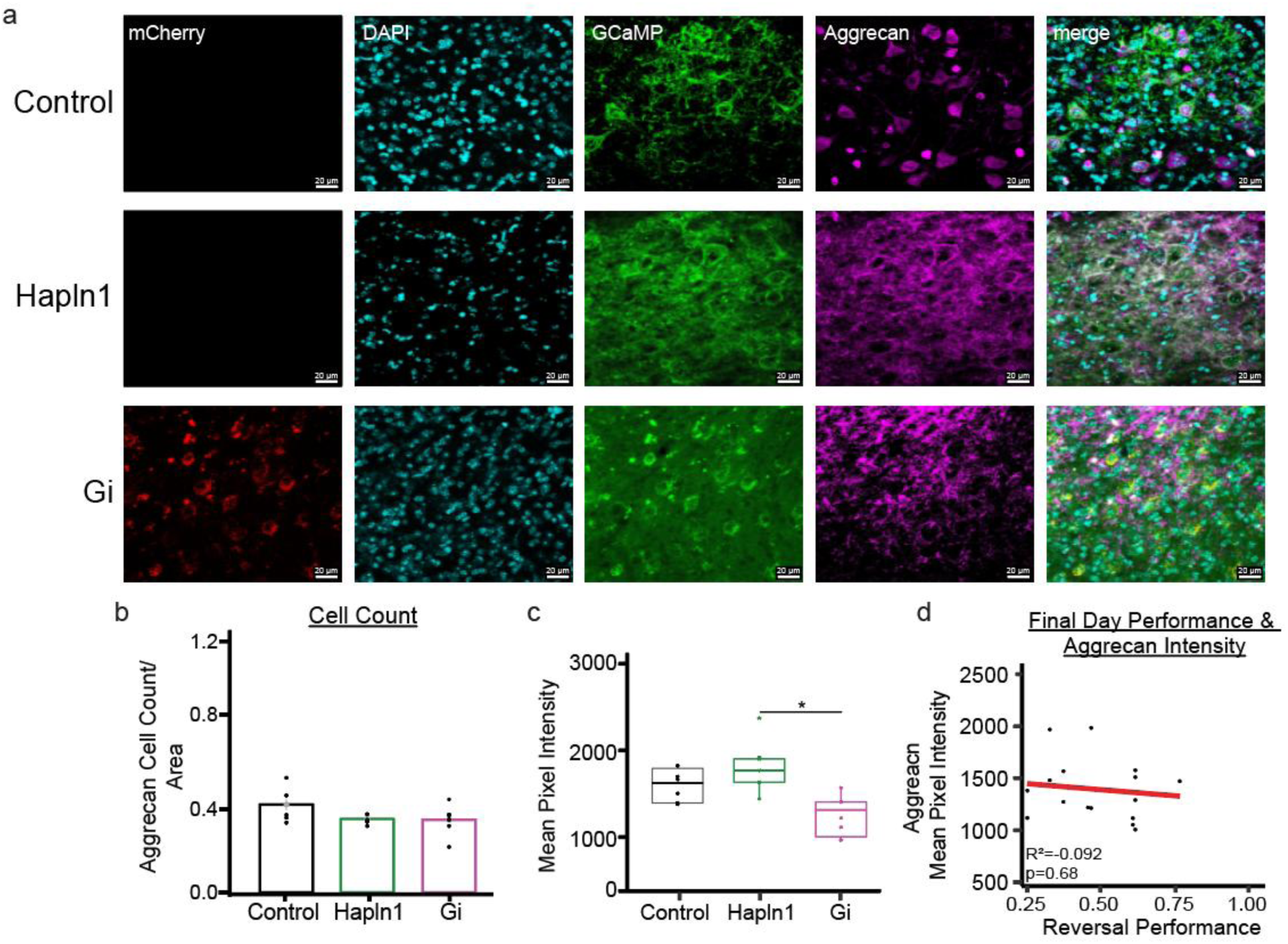
Adolescent LCN perturbation alters aggrecan intensity between experimental conditions. (A.) Immunofluorescence shows Gi DREADDs (mCherry), GCaMP (green), and aggrecan positive cells stained for anti-aggrecan (magenta) in the lateral cerebellar nucleus (LCN) in Control, Hapln1, and Gi animals (Scale bar: 20 μm, taken at 40x objective). (B.) No differences were found in aggrecan cell count between groups (F(2,35)=1.09, p=0.35, ANOVA). (C.) Analysis of LCN aggrecan intensity showed a group difference (F(2,35)=3.76, p=0.03, AVNOA) driven by Hapln1 animals (p=0.03, Tukey) compared to Gi animals. (D.) LCN Aggrecan and reversal performance. (A.) Pearson correlation between aggrecan intensity and reversal performance. There was no significant relationship identified between aggrecan and reversal performance (R²=-0.092 p=0.68). Data represents mean ± SEM, n=6 mice per group, *p < 0.05.

## References

1. Ito, M. (2002). Historical review of the significance of the cerebellum and the role of Purkinje cells in motor learning. Ann. N. Y. Acad. Sci. 978, 273–288. 10.1111/j.1749-6632.2002.tb07574.x.

2. Lyle, T.T., and Verpeut, J.L. (2025). Adolescent cerebellar nuclei manipulation alters reversal learning and perineuronal net intensity independently in male and female mice. J. Neurosci., e2182232024. 10.1523/JNEUROSCI.2182-23.2024.

3. Lyle, T.T., Masho Elbeh, K., Chambers, D., Vieira, H., and Verpeut, J.L. (2025). Attenuation of social preference and alteration of cortical neurons following adolescent cerebellar nuclei perturbation. Mol. Psychiatry, 1–15. 10.1038/s41380-025-03404-3.

4. Verpeut, J.L., and Oostland, M. (2024). The significance of cerebellar contributions in early-life through aging. Front. Comput. Neurosci. 18. 10.3389/fncom.2024.1449364.

5. Verpeut, J.L., Bergeler, S., Kislin, M., William Townes, F., Klibaite, U., Dhanerawala, Z.M., Hoag, A., Janarthanan, S., Jung, C., Lee, J., et al. (2023). Cerebellar contributions to a brainwide network for flexible behavior in mice. Commun Biol 6, 605. 10.1038/s42003-023-04920-0.

6. Wang, S.S.-H., Kloth, A.D., and Badura, A. (2014). The cerebellum, sensitive periods, and autism. Neuron 83, 518–532. 10.1016/j.neuron.2014.07.016.

7. Badura, A., Verpeut, J.L., Metzger, J.W., Pereira, T.D., Pisano, T.J., Deverett, B., Bakshinskaya, D.E., and Wang, S.S.-H. (2018). Normal cognitive and social development require posterior cerebellar activity. Elife 7. 10.7554/eLife.36401.

8. Stoodley, C.J., and Schmahmann, J.D. (2009). The cerebellum and language: evidence from patients with cerebellar degeneration. Brain Lang. 110, 149–153. 10.1016/j.bandl.2009.07.006.

9. Kim, L.H., Heck, D.H., and Sillitoe, R.V. (2024). Cerebellar functions beyond movement and learning. Annu. Rev. Neurosci. 47, 145–166. 10.1146/annurev-neuro-100423-104943.

10. Middleton, F.A., and Strick, P.L. (2001). Cerebellar projections to the prefrontal cortex of the primate. J. Neurosci. 21, 700–712. 10.1523/jneurosci.21-02-00700.2001.

11. Kelly, R.M., and Strick, P.L. (2003). Cerebellar loops with motor cortex and prefrontal cortex of a nonhuman primate. J. Neurosci. 23, 8432–8444. 10.1523/jneurosci.23-23-08432.2003.

12. E, K.-H., Chen, S.-H.A., Ho, M.-H.R., and Desmond, J.E. (2014). A meta-analysis of cerebellar contributions to higher cognition from PET and fMRI studies. Hum. Brain Mapp. 35, 593–615. 10.1002/hbm.22194.

13. Pisano, T.J., Dhanerawala, Z.M., Kislin, M., Bakshinskaya, D., Engel, E.A., Hansen, E.J., Hoag, A.T., Lee, J., de Oude, N.L., Venkataraju, K.U., et al. (2021). Homologous organization of cerebellar pathways to sensory, motor, and associative forebrain. Cell Rep. 36, 109721. 10.1016/j.celrep.2021.109721.

14. Nguyen, T.M., Thomas, L.A., Rhoades, J.L., Ricchi, I., Yuan, X.C., Sheridan, A., Hildebrand, D.G.C., Funke, J., Regehr, W.G., and Lee, W.-C.A. (2023). Structured cerebellar connectivity supports resilient pattern separation. Nature 613, 543–549. 10.1038/s41586-022-05471-w.

15. Arleo, A., Bareš, M., Bernard, J.A., Bogoian, H.R., Bruchhage, M.M.K., Bryant, P., Carlson, E.S., Chan, C.C.H., Chen, L.-K., Chung, C.-P., et al. (2024). Consensus paper: Cerebellum and ageing. Cerebellum 23, 802–832. 10.1007/s12311-023-01577-7.

16. Ahmadian, N., van Baarsen, K., van Zandvoort, M., and Robe, P.A. (2019). The cerebellar cognitive affective syndrome-a meta-analysis. Cerebellum 18, 941–950. 10.1007/s12311-019-01060-2.

17. Schmahmann, J.D., Weilburg, J.B., and Sherman, J.C. (2007). The neuropsychiatry of the cerebellum - insights from the clinic. Cerebellum 6, 254–267. 10.1080/14734220701490995.

18. Ranjbar, H., Soti, M., Banazadeh, M., Saleki, K., Kohlmeier, K.A., and Shabani, M. (2021). Addiction and the cerebellum with a focus on actions of opioid receptors. Neurosci. Biobehav. Rev. 131, 229–247. 10.1016/j.neubiorev.2021.09.021.

19. Darvishzadeh-Mahani, F., Rajabi, S., Alehashem, M., Alaei, H., and Ramshini, E. (2025). Psychostimulants and the cerebellum: Does the cerebellum get involved in the abuse of methamphetamine and cocaine? Prog. Neuropsychopharmacol. Biol. Psychiatry 142, 111479. 10.1016/j.pnpbp.2025.111479.

20. D’Mello, A.M., and Stoodley, C.J. (2015). Cerebro-cerebellar circuits in autism spectrum disorder. Front. Neurosci. 9, 408. 10.3389/fnins.2015.00408.

21. Olivito, G., Clausi, S., Laghi, F., Tedesco, A.M., Baiocco, R., Mastropasqua, C., Molinari, M., Cercignani, M., Bozzali, M., and Leggio, M. (2017). Resting-State Functional Connectivity Changes Between Dentate Nucleus and Cortical Social Brain Regions in Autism Spectrum Disorders. Cerebellum 16, 283–292. 10.1007/s12311-016-0795-8.

22. LeBlanc, J.J., and Fagiolini, M. (2011). Autism: a “critical period” disorder? Neural Plast. 2011, 921680. 10.1155/2011/921680.

23. del Río, J.A., de Lecea, L., Ferrer, I., and Soriano, E. (1994). The development of parvalbumin-immunoreactivity in the neocortex of the mouse. Brain Res. Dev. Brain Res. 81, 247–259. 10.1016/0165-3806(94)90311-5.

24. Chattopadhyaya, B., Di Cristo, G., Higashiyama, H., Knott, G.W., Kuhlman, S.J., Welker, E., and Huang, Z.J. (2004). Experience and activity-dependent maturation of perisomatic GABAergic innervation in primary visual cortex during a postnatal critical period. J. Neurosci. 24, 9598–9611. 10.1523/JNEUROSCI.1851-04.2004.

25. Hu, H., Gan, J., and Jonas, P. (2014). Interneurons. Fast-spiking, parvalbumin^+^ GABAergic interneurons: from cellular design to microcircuit function. Science 345, 1255263. 10.1126/science.1255263.

26. Bartos, M., Vida, I., and Jonas, P. (2007). Synaptic mechanisms of synchronized gamma oscillations in inhibitory interneuron networks. Nat. Rev. Neurosci. 8, 45–56. 10.1038/nrn2044.

27. Uusisaari, M., Obata, K., and Knöpfel, T. (2007). Morphological and electrophysiological properties of GABAergic and non-GABAergic cells in the deep cerebellar nuclei. J. Neurophysiol. 97, 901–911. 10.1152/jn.00974.2006.

28. Udakis, M., Pedrosa, V., Chamberlain, S.E.L., Clopath, C., and Mellor, J.R. (2020). Interneuron-specific plasticity at parvalbumin and somatostatin inhibitory synapses onto CA1 pyramidal neurons shapes hippocampal output. Nat. Commun. 11, 4395. 10.1038/s41467-020-18074-8.

29. Nissen, W., Szabo, A., Somogyi, J., Somogyi, P., and Lamsa, K.P. (2010). Cell type-specific long-term plasticity at glutamatergic synapses onto hippocampal interneurons expressing either parvalbumin or CB1 cannabinoid receptor. J. Neurosci. 30, 1337–1347. 10.1523/JNEUROSCI.3481-09.2010.

30. Vickers, E.D., Clark, C., Osypenko, D., Fratzl, A., Kochubey, O., Bettler, B., and Schneggenburger, R. (2018). Parvalbumin-interneuron output synapses show spike-timing-dependent plasticity that contributes to auditory map remodeling. Neuron 99, 720–735.e6. 10.1016/j.neuron.2018.07.018.

31. Fawcett, J.W., Oohashi, T., and Pizzorusso, T. (2019). The roles of perineuronal nets and the perinodal extracellular matrix in neuronal function. Nat. Rev. Neurosci. 20, 451–465. 10.1038/s41583-019-0196-3.

32. Sanchez, B., Kraszewski, P., Lee, S., and Cope, E.C. (2024). From molecules to behavior: Implications for perineuronal net remodeling in learning and memory. J. Neurochem. 168, 1854–1876. 10.1111/jnc.16036.

33. Carulli, D., Broersen, R., de Winter, F., Muir, E.M., Mešković, M., de Waal, M., de Vries, S., Boele, H.-J., Canto, C.B., De Zeeuw, C.I., et al. (2020). Cerebellar plasticity and associative memories are controlled by perineuronal nets. Proc. Natl. Acad. Sci. U. S. A. 117, 6855– 6865. 10.1073/pnas.1916163117.

34. Romberg, C., Yang, S., Melani, R., Andrews, M.R., Horner, A.E., Spillantini, M.G., Bussey, T.J., Fawcett, J.W., Pizzorusso, T., and Saksida, L.M. (2013). Depletion of perineuronal nets enhances recognition memory and long-term depression in the perirhinal cortex. J. Neurosci. 33, 7057–7065. 10.1523/JNEUROSCI.6267-11.2013.

35. Dauth, S., Grevesse, T., Pantazopoulos, H., Campbell, P.H., Maoz, B.M., Berretta, S., and Parker, K.K. (2016). Extracellular matrix protein expression is brain region dependent: Brain Extracellular Matrix Distribution Analysis. J. Comp. Neurol. 524, 1309–1336. 10.1002/cne.23965.

36. Rowlands, D., Lensjø, K.K., Dinh, T., Yang, S., Andrews, M.R., Hafting, T., Fyhn, M., Fawcett, J.W., and Dick, G. (2018). Aggrecan Directs Extracellular Matrix-Mediated Neuronal Plasticity. J. Neurosci. 38, 10102–10113. 10.1523/JNEUROSCI.1122-18.2018.

37. Shaw, K.A., Williams, S., Patrick, M.E., Valencia-Prado, M., Durkin, M.S., Howerton, E.M., Ladd-Acosta, C.M., Pas, E.T., Bakian, A.V., Bartholomew, P., et al. (2025). Prevalence and early identification of autism spectrum disorder among children aged 4 and 8 years -autism and Developmental Disabilities Monitoring Network, 16 sites, United States, 2022. MMWR Surveill. Summ. 74, 1–22. 10.15585/mmwr.ss7402a1.

38. Rhodes, M.E., and Frye, C.A. (2004). Estrogen has mnemonic-enhancing effects in the inhibitory avoidance task. Pharmacol. Biochem. Behav. 78, 551–558. 10.1016/j.pbb.2004.03.025.

39. Walf, A.A., Koonce, C., Manley, K., and Frye, C.A. (2009). Proestrous compared to diestrous wildtype, but not estrogen receptor beta knockout, mice have better performance in the spontaneous alternation and object recognition tasks and reduced anxiety-like behavior in the elevated plus and mirror maze. Behav. Brain Res. 196, 254–260. 10.1016/j.bbr.2008.09.016.

40. Bankhead, P., Loughrey, M.B., Fernández, J.A., Dombrowski, Y., McArt, D.G., Dunne, P.D., McQuaid, S., Gray, R.T., Murray, L.J., Coleman, H.G., et al. (2017). QuPath: Open source software for digital pathology image analysis. Sci. Rep. 7, 16878. 10.1038/s41598-017-17204-5.

41. Simpson, E.H., Akam, T., Patriarchi, T., Blanco-Pozo, M., Burgeno, L.M., Mohebi, A., Cragg, S.J., and Walton, M.E. (2024). Lights, fiber, action! A primer on in vivo fiber photometry. Neuron 112, 718–739. 10.1016/j.neuron.2023.11.016.

42. Bruno, C.A., O’Brien, C., Bryant, S., Mejaes, J.I., Estrin, D.J., Pizzano, C., and Barker, D.J. (2021). pMAT: An open-source software suite for the analysis of fiber photometry data. Pharmacol. Biochem. Behav. 201, 173093. 10.1016/j.pbb.2020.173093.

43. Darmohray, D.M., Jacobs, J.R., Marques, H.G., and Carey, M.R. (2019). Spatial and Temporal Locomotor Learning in Mouse Cerebellum. Neuron 102, 217–231.e4. 10.1016/j.neuron.2019.01.038.

44. Kim, C.H., Hvoslef-Eide, M., Nilsson, S.R.O., Johnson, M.R., Herbert, B.R., Robbins, T.W., Saksida, L.M., Bussey, T.J., and Mar, A.C. (2016). Erratum to: The continuous performance test (rCPT) for mice: a novel operant touchscreen test of attentional function. Psychopharmacology 233, 3471. 10.1007/s00213-016-4400-0.

45. Suttkus, A., Rohn, S., Weigel, S., Glöckner, P., Arendt, T., and Morawski, M. (2014). Aggrecan, link protein and tenascin-R are essential components of the perineuronal net to protect neurons against iron-induced oxidative stress. Cell Death Dis. 5, e1119. 10.1038/cddis.2014.25.

46. Otsuka, M.Y., Essel, L.B., Sinha, A., Nickerson, G., Mejia, S.M., Matthews, R.T., and Bouyain, S. (2024). The hyaluronan-binding activity of aggrecan is important, but not essential, for its specific insertion into perineuronal nets. bioRxiv. 10.1101/2024.11.25.625086.

47. Favuzzi, E., Marques-Smith, A., Deogracias, R., Winterflood, C.M., Sánchez-Aguilera, A., Mantoan, L., Maeso, P., Fernandes, C., Ewers, H., and Rico, B. (2017). Activity-Dependent Gating of Parvalbumin Interneuron Function by the Perineuronal Net Protein Brevican. Neuron 95, 639–655.e10. 10.1016/j.neuron.2017.06.028.

48. Galtrey, C.M., and Fawcett, J.W. (2007). The role of chondroitin sulfate proteoglycans in regeneration and plasticity in the central nervous system. Brain Res. Rev. 54, 1–18. 10.1016/j.brainresrev.2006.09.006.

49. Miyata, S., and Kitagawa, H. (2016). Chondroitin 6-sulfation regulates perineuronal net formation by controlling the stability of aggrecan. Neural Plast. 2016, 1305801. 10.1155/2016/1305801.

50. Brain Knowledge Platform https://knowledge.brain-map.org/data/LVDBJAW8BI5YSS1QUBG/explore?filterOptions.

